# Transcriptional downregulation of rhodopsin is associated with desensitization of rods to light-induced damage in a murine model of retinitis pigmentosa

**DOI:** 10.1101/2025.04.03.646684

**Authors:** Shimpei Takita, Hemavathy Harikrishnan, Masaru Miyagi, Yoshikazu Imanishi

## Abstract

Class I rhodopsin mutations are known for some of the most severe forms of vision impairments in dominantly inherited rhodopsin retinitis pigmentosa. They disrupt the VxPx transport signal, which is required for the proper localization of rhodopsin to the outer segments. While various studies have focused on the light-dependent toxicity of mutant rhodopsin, it remains unclear whether and how these mutations exert dominant-negative effects. Using the class I *Rho^Q344X^* rhodopsin knock-in mouse model, we characterized the expression of rhodopsin and other genes by RNA sequencing and qPCR. Those studies indicated that rhodopsin is the most prominently downregulated photoreceptor-specific gene in *Rho^Q344X/+^* mice. Rhodopsin is downregulated significantly prior to the onset of rod degeneration, whereas downregulation of other phototransduction genes, transducin*α*, and Pde6α, occurs after the onset and correlate with the degree of rod cell loss. Those studies indicated that the mutant rhodopsin gene causes downregulation of wild-type rhodopsin, imposing an mRNA-level dominant negative effect. Moreover, it causes downregulation of the mutant mRNA itself, mitigating the toxicity. The observed dominant effect is likely common among rhodopsin retinitis pigmentosa as we found a similar rhodopsin downregulation in the major class II rhodopsin mutant model, *Rho^P23H/+^* mice, in which mutant rhodopsin is prone to misfold. Potentially due to mitigated toxicity by reduced rhodopsin expression, *Rho^Q344X/+^* mice did not exhibit light-dependent exacerbation of rod degeneration, even after continuous exposure of mice for 5 days at 3000 lux. Thus, this study describes a novel form of dominant negative effect in inherited neurodegenerative disorders.

## Introduction

Mutations in the rhodopsin gene (*Rho*) are the primary causes of autosomal dominant retinitis pigmentosa (adRP) (1). Two types of mutations have been mainly studied in the past, one categorized as class I and the other as class II (1). Class I mutations, exemplified by *Rho^Q344X^*, disrupt the VxPx trafficking signal located at the C-terminal tail of rhodopsin. Class II mutations, exemplified by *Rho^P23H^*, cause opsin misfolding and are the most prevalent type of defects found among adRP patients. The dominant inheritance pattern is possibly attributable to a combination of toxic gain-of-function and other effects of the mutant alleles. Supporting the toxic gain-of-function role, these mutations can cause rhodopsin to misfold, become mislocalized, or exhibit abnormal signaling behaviors, ultimately compromising the function and survival of photoreceptor cells (1). The gain-of-function role of mutations is also indicated phenotypically; homozygous adRP rhodopsin mutations cause rod degeneration much more rapidly than homozygous loss-of-function mutations (2–4). Less studied are the dominant-negative effects of mutant rhodopsin, which lead to loss of function and compromised rod photoreceptor activity. Rhodopsin molecules are prone to dimerization (5, 6).

Through heterodimerization, mutant rhodopsins are suggested to impair the function of wild-type rhodopsin, exerting dominant-negative effects at the protein level (7). Consequently, the structural or localization anomalies observed in mutant rhodopsin may propagate to wild-type rhodopsin molecules by inducing co-aggregation or co-mislocalization. Despite these structural implications, *in vivo* evidence of such propagation has not been sufficient to establish whether mutated rhodopsin proteins impose a dominant-negative effect on wild-type rhodopsin, potentially further compromising the functionality of rod photoreceptor cells. Although the possible dimerization of rhodopsin molecules has been suggested, the mislocalization of wild-type rhodopsin was not observed in transgenic *Xenopus laevis* expressing the *Rho^Q344X^* mutant of either *Xenopus* (8) or human origin (9). In the knock-in mouse model with the *Rho^Q344X^* mutation (4), wild-type rhodopsin is mislocalized, though to a much lesser degree than the mutant rhodopsin, suggesting dimerization-based dominant negative effects, if any exist, are minimal. However, we do observe a pronounced reduction in wild-type rhodopsin protein levels, the mechanisms of which remain uncharacterized (4). Clarifying how these loss-of-function mechanisms interact with well-established gain-of-function effects is essential for fully understanding disease pathogenesis and clinical manifestations in rhodopsin adRP.

Light is considered one of the major environmental factors contributing to rod degeneration caused by rhodopsin gene mutations (10–13). In healthy photoreceptor cells, light triggers activation of the phototransduction cascade, which is confined to the outer segment (OS). This compartmentalization is crucial for the rapid and accurate transmission of biochemical and electrical signals in photoreceptors (14). This compartmentalization is disrupted in the case of rhodopsin mislocalization, where light triggers ectopic activation of phototransduction and other G protein-mediated pathways. Insights into the effects of disrupted compartmentalization initially came from studies on salamander rod photoreceptors cultured for extended periods, during which rhodopsin mislocates across the entire plasma membrane (15). In this and a transgenic zebrafish model of class I rhodopsin mutation, activation of rhodopsin in non-OS compartments, such as the inner segments (IS), cell body, and synapse, may increase adenylate cyclase activity, leading to cAMP synthesis (15, 16). This ectopic activation of rhodopsin can worsen rod photoreceptor death. Research in transgenic mouse models with class I rhodopsin mutations supports these findings (13). The degree to which light can exacerbate rod degeneration likely depends on the extent of rhodopsin mislocalization. However, a significant limitation of earlier studies is their reliance on transgenic models and other experimental systems that overexpress trafficking-deficient rhodopsin or induce its elevated mislocalization, resulting in non-physiological conditions that may not accurately reflect the disease mechanisms in patients. These approaches do not accurately reflect the specific level of rhodopsin mislocalization found in adRP, which is crucial for understanding the condition’s pathophysiology. The majority of patients with rhodopsin mutations are heterozygous carriers, meaning their photoreceptor cells produce both mutated and wild-type proteins at similar transcriptional levels as seen for the knock-in mouse models of adRP (3, 4).

In the present study using the *Rho^Q344X^* knock-in mouse (4), we revisited the dominant negative effect of the *Rho^Q344X^* on the wild-type rhodopsin. Dominant negative effect would result in loss of rhodopsin function, reduced photoreceptor sensitivity and compromised cell survival, as demonstrated in rhodopsin knockout mice (2). Employing high-throughput proteomics and RNA sequencing (RNA-seq) technologies, we unexpectedly found the dominant negative effect to be at the transcriptional level, likely attenuating the function of wild-type rhodopsin while mitigating the toxicity of mislocalized rhodopsin. As this model expresses the adequate level of mutant rhodopsin, we revisited the effects of light on the degeneration and function of rod photoreceptors. Under physiological cyclic light conditions, light did not exacerbate rod degeneration, suggesting ectopic rhodopsin activation, if any occurs, has minimal impact on the survival of rods. Moreover, light enhanced the dominant effect which acts in a negative feedback loop to further mitigate rhodopsin toxicity. Thus, our studies of the rhodopsin-RP model reveal novel inter-allelic interactions impacting the pathophysiology of the devastating blinding disorders.

## Results

### The *Rho^Q344X^* allele exerts a dominant-negative effect on the *Rho* allele, downregulating both Q344X and wild-type *Rho* mRNA expressions

To comprehensively understand and compare the transcriptome profiles of *Rho^Q344X/+^* and wild-type mice, we conducted deep RNA-seq of their retinas at postnatal days (P) 35, a time point at which a substantial fraction of rods (∼80%) remains viable in *Rho^Q344X/+^* mice (Figure 1 and Supplemental Table S1) (4). Mice were reared under standard housing condition of cyclic 12 hours light (∼200 lux) and 12 hours dark. In wild-type retinas, *Rho* was the second most abundantly expressed gene following the mitochondrial cytochrome c oxidase I (mt-Co1) gene (Figure 1A). In the *Rho^Q344X/+^*knock-in retinas, we noted significant downregulation of *Rho* mRNA expression (Figures 1A), based on analyses of individual genes (Supplemental Table S2). Retinal expression of *Rho* was approximately three times lower in *Rho^Q344X/+^*than in wild-type mice. We conducted a similar analysis for P35 *Rho^P23H/+^*mice (Supplemental Figure S1), a time point at which ∼70% of rods remain (3, 4), and found four-fold downregulation of *Rho* mRNA in this genotype (5255.75 for wild-type, 1828.76 for *Rho^Q344X/+^*, and 1299.00 for *Rho^P23H/+^* in FPKM [Fragments Per Kilobase of exon per Million mapped reads]). Quantitative analyses indicated that the expression levels of most photoreceptor-specific transcripts declined less than 50% as a result of the Q344X mutation. Photoreceptor-specific transcripts such as *Gnat1* (G protein subunit alpha transducin 1, 34.1% decline from wild-type) and Peripherin-2 (*Prph2*, 22.0% decline from wild-type) showed less than 50% decline in their expression levels. Those declines are correlated with a 20–25% loss of rod photoreceptor cells by P35 (4). Unlike other photoreceptor-specific genes, we found a disproportionately high degree of loss in *Rho* gene transcripts (by 65-70%) in *Rho^Q344X/+^* and *Rho^P23H/+^* mice, suggesting this specific downregulation is a characteristic of rhodopsin adRP models.

**Figure 1.**
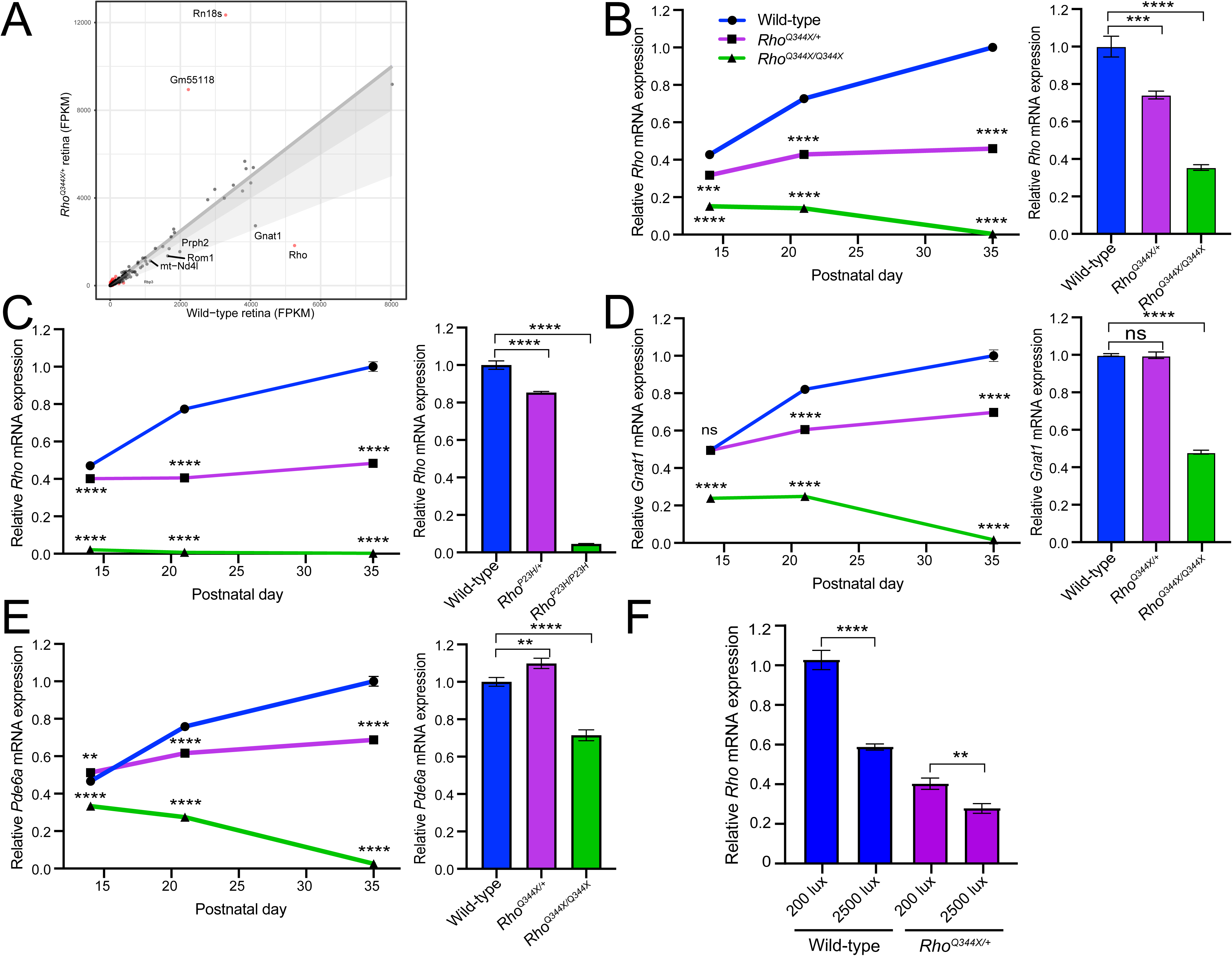
*Rhodopsin* (*Rho*) mRNA is downregulated prior to the onset of photoreceptor degeneration in the *Rho^Q344X/+^* and *Rho^P23H/+^* retinas. (A) RNA expression levels of individual genes were plotted to compare between P35 wild-type and *Rho^Q344X/+^* mouse retinas reared under standard cyclic 200 lux condition (x- and y-axes, respectively). A total of 851 genes (shown in red dots) were differentially expressed with 2-fold change (log2-transformed change of < -1 or > 1) and q-value of < 0.05 thresholds. *Rhodopsin* was the second most abundantly expressed gene in wild-type retinas and was downregulated by 70% in *Rho^Q344X/+^* retinas. The dark gray shade indicates less than a 20% decline in mRNA expressions. The light gray shade indicates less than a 50% decline in mRNA expressions. (B) *Rho* mRNA expression levels were measured by quantitative PCR (qPCR) at P14, P21, and P35 using wild-type (blue), *Rho^Q344X/+^*(purple), and *Rho^Q344X/Q344X^* (green) retinas reared under standard cyclic 200 lux light condition. The temporal *Rho* mRNA expression levels were plotted as a function of postnatal days (P14 – 35), and the data are presented as mean ± SD (n = 3 animals for each genotype). (C) *Rho* mRNA expression levels were quantitatively measured by qPCR at P14, P21, and P35 using wild-type (blue), *Rho^P23H/+^*(purple), and *Rho^P23H/P23H^* (green) mouse retinas reared under standard cyclic 200 lux light condition. The temporal *Rho* mRNA expression levels were plotted as a function of postnatal days (P14 – 35). (D and E) *Gnat1* (D) or *Pde6a* (E) mRNA expression levels were measured by qPCR at P14, P21, and P35 using wild-type (blue), *Rho^Q344X/+^* (purple), and *Rho^Q344X/Q344X^*(green) retinas reared under standard cyclic 200 lux light condition. The temporal mRNA expression levels were plotted as a function of postnatal days (P14 – 35). (F) *Rho* mRNA expression levels were measured by qPCR at P21 using wild-type (blue) and *Rho^Q344X/+^* (purple) retinas reared under either 200 lux or 2500 lux cyclic light conditions. The data are represented as mean ± SD (n = 3 animals for each genotype). The data were subjected to statistical analysis using two-way ANOVA, followed by Tukey’s post-hoc test for pairwise comparisons. Statistical significance (****, *p* < 0.0001; ***, *p* < 0.001, ns, not significant), comparing *Rho^Q344X/+^* vs wild-type or *Rho^Q344X/Q344X^* vs wild-type, is indicated above each data point.

To verify and further investigate whether the observed downregulation of *Rho* mRNA exceeds the degree of rod photoreceptor loss, we quantified mRNA levels at P14, P21, and P35, the time course corresponding to before, at the onset (4), and soon after the onset of photoreceptor degeneration (Figure 1B, left panel) in *Rho^Q344X/+^* mice using quantitative PCR (qPCR). Unexpectedly, downregulation of *Rho* mRNA was observed prior to the onset of rod degeneration (P14; Figure 1B, right panel), and thus loss of rod photoreceptor is not the primary reason for the observed downregulation. In *Rho^Q344X/+^* mice, *Rho* mRNA levels were approximately 74% of those observed in wild-type mice at P14 and about 59% at P21. In *Rho^P23H/+^* mice (Figure 1C), *Rho* mRNA levels were approximately 85% at P14 (Figure 1C, right panel) and 52% at P21. In both *Rho^Q344X/+^* and *Rho^P23H/+^*mice, the degree of *Rho* mRNA loss is more pronounced than the extent of rod loss, which has not occurred at P14 and is barely seen at P21 (4, 17). At P35, *Rho* levels were reduced by 54% in *Rho^Q344X/+^* mice and by 51.7% in *Rho^P23H/+^*mice. Given that the wild-type and mutant *Rho* alleles are expressed at approximately equal levels in both genotypes (3, 4), the observed over 50% reduction indicates that both the wild-type and mutant alleles undergo downregulation in rhodopsin adRP models.

In *Rho^Q344X/+^* mice, other rod photoreceptor-specific transcripts, such as *Gnat1* and *Pde6a* (phosphodiesterase 6A, cGMP-specific, rod, alpha encoding the alpha subunit of PDE6), did not show downregulation at P14 (Figure 1D and E, right panels). Their downregulation became observable at P21 and was more pronounced by P35 (Figure 1D and E, left panels). The downregulation of these rod-specific transcripts was approximately 30% at P35, aligning with the extent of rod photoreceptor cell loss observed in *Rho^Q344X/+^* mice at this stage (4).

*Rho^Q344X/Q344X^* and *Rho^P23H/P23H^* homozygous mice exhibited marked *Rho* mRNA downregulation at P14 (Figure 1B and C, right panels), with expression levels reduced to 35.4% and 4.6% of wild-type levels, respectively. Since rod degeneration begins before P14 in these genotypes, the observed downregulation is at least partially attributable to the rapid loss of rod photoreceptor cells. To further assess the contribution of degeneration in *Rho^Q344X/Q344X^* mice, we analyzed the expression of *Gnat1* and *Pde6a*. Both genes showed significant downregulation at P14, with expression levels reduced to 48.1% and 71.5% of wild-type levels, respectively (Figure 1D and E, right panels). Interestingly, the degrees of downregulation for *Gnat1* and *Pde6a* were less pronounced than that observed for *Rho*, suggesting that *Rho* downregulation in these rhodopsin mutant models is not solely due to rod photoreceptor cell loss.

The mRNA levels of *Rho*, *Gnat1*, and *Pde6a* progressively declined between P14 and P35, reflecting the near-complete loss of rod photoreceptors in *Rho^Q344X/Q344X^*mice during this period.

### Exacerbation of rod degeneration by light exposure is minimal in the *Rho^Q344X/+^* mice

Previous studies indicated that rod degeneration can be attributed to the light-activated mislocalized rhodopsin in IS and other non-OS compartments of rod (13, 15, 16). Those studies were conducted using mouse and zebrafish transgenic animals where class I mutant rhodopsin is overexpressed (13, 16) or cultured and dissociated salamander rod cells in which rhodopsin mislocalization is observed (15). Given that the light effect is likely to depend on the amount of ectopically light-activated rhodopsin due to mislocalization, our *Rho^Q344X/+^* knock-in mouse model demonstrating appropriate levels of Rho^Q344X^ expression, with equal amounts of mutant and wild-type *Rho* alleles expressed, is ideal to accurately mimic the physiological condition of adRP.

To test the effects of light exposure, we employed two different intensities at physiologically relevant levels: 200 lux and 2500 lux (18–20). *Rho^Q344X/+^*mice were reared under a cycle of 12 hours of light (200 lux or 2500 lux) and 12 hours of darkness (12 h L/D cycles) or under continuous darkness (24 h darkness). The thickness of the outer nuclear layer (ONL), where photoreceptor nuclei are located, was measured using optical coherence tomography (OCT) and compared among cyclic light-reared and dark-reared mice from P21 to P120 (Figure 2A–C). ONL thickness in cyclic light-reared animals did not significantly differ from that of dark-reared animals from P35 to P120 (Figure 2D). We found no significant decrease in ONL thicknesses due to light exposure, at either 200 lux or 2500 lux. In all the quadrants of the retina, ONL thicknesses were similar among all the cohorts of animals, indicative of an absence of light-exposure effects, if they existed, should have resulted in significantly thinner ONL in the ventral retina which is exposed to more intense light than other quadrants. At P21, however, dark-reared animals showed slightly thinner ONL, with variable degrees. It is unlikely that darkness promoted the degeneration of photoreceptors as ONL thicknesses are similar at subsequent stages of P35 – 120, regardless of light conditions. Collectively, our experiments indicate that increasing light stimuli under physiological conditions (200 lux and 2500 lux) does not have a major impact on the degree of photoreceptor degeneration, thus, indicating ectopic rhodopsin light-activation does not contribute to rod degeneration in *Rho^Q344X/+^*animals. This lack of light effect is consistent with the observation that the expression of rhodopsin, a chromophore critical for light-mediated degeneration (15, 20), is significantly downregulated at the mRNA level (Figure 1F) and protein level (4) in *Rho^Q344X/+^*mice reared under 200 lux cyclic light. By qPCR, we found that exposure of mice to 2500 lux resulted in a further reduction of *Rho* mRNA levels by 58.8% in wild-type and by 31.0% in *Rho^Q344X/+^* mice (Figure 1F). These results suggest that *Rho* downregulation in *Rho^Q344X/+^* mice is neuroprotective, as it helps mitigate the rhodopsin phototoxicity.

**Figure 2.**
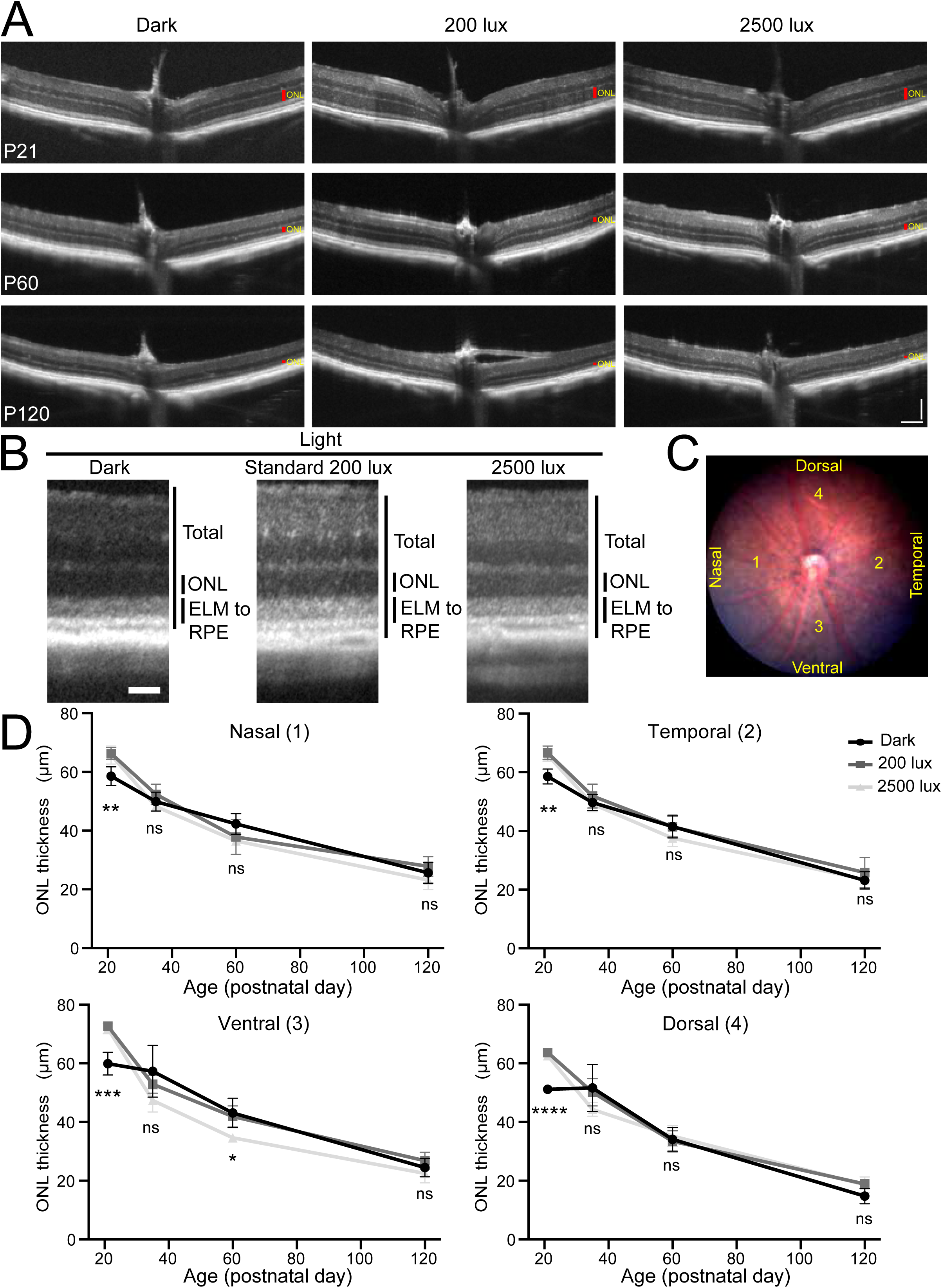
OCT analyses of class I *Rho^Q344X^* knock-in mutant mouse indicate similar degeneration of photoreceptors under three light intensities. (A) OCT images spanning the ventral-optic nerve head (ONH)-dorsal regions were acquired from *Rho^Q344X/+^* mice at P21, P60, and P120. ONLs are indicated by red vertical bars. Scale bars, 100 µm. (B) OCT images were acquired at locations 500 µm away from the ONH from P60 *Rho^Q344X/+^* mice reared under dark, 200 lux, or 2500 lux light conditions. ELM, external limiting membrane; RPE, retinal pigment epithelium. Scale bar, 50 µm. (C) A representative fundus image of P60 *Rho^Q344X/+^* mice is shown. The thicknesses of the ONL were measured at four distinct locations situated 500 µm away from the ONH, as illustrated in the fundus image of *Rho^Q344X/+^*mice reared under dark condition. (D) ONL thicknesses were compared among dark (black circle), 200 lux (dark gray rectangle), and 2500 lux (light gray triangle) light conditions in four distinct regions (nasal, temporal, ventral, and dorsal) of *Rho^Q344X/+^* retinas as indicated in (C). The thicknesses of the ONL were plotted as a function of postnatal days (P21 – 120). The data are represented as mean ± SD (n = 4 animals for each genotype). The data were subjected to statistical analysis using two-way ANOVA, followed by Dunnett’s multiple comparisons test for pairwise comparisons. Statistical significance (****, *p* < 0.0001; ***, *p* < 0.001; \**p* < 0.01; ns, not significant) is shown below each data point (dark vs 200 lux or dark vs 2500 lux).

To further understand molecular changes induced by light exposure, we compared the retinal transcriptomic data from dark-reared and standard 200-lux-cyclic-light-reared animals (Figure 3). Principal component analysis (PCA) revealed that *Rho^Q344X/+^* mice displayed a significant divergence from wild-type mice, independent of the light conditions (Figure 3A and B). This was reflected by Pearson correlation coefficients of 0.62 under dark condition and 0.87 under standard cyclic 200 lux light condition. The transcriptome profiles of *Rho^Q344X/+^* mice reared under dark and standard cyclic 200 lux light conditions were similar, with a Pearson correlation of 0.95. In contrast, wild-type mice exhibited notable transcriptomic differences between the two light conditions, with a Pearson correlation of 0.45. These results suggest that light exposure has minimal impact on the transcriptome, consistent with the absence of light-dependent rod degeneration in *Rho^Q344X/+^*mice. Through pathway analysis employing gene ontology (GO) terms (Supplemental Figure S2) identified that genes involved in inflammation-related pathways such as regulation of adaptive immune response (GO0002819), response to type II interferon (GO0034341), and regulation of tumor necrosis factor production (GO0032680) such as Ifitm proteins, Ifitm1-3 (interferon induced transmembrane protein 1-3, 264.8%, 244.1%, and 415.1% wild-type, respectively), B2m (beta-2-microglobulin, 312.1% wild-type) and C3 (complement C3, 372.5% wild-type) were upregulated reflecting the rod photoreceptor degeneration caused by the *Rho^Q344X^*mutation (Supplemental Figure S3).

**Figure 3.**
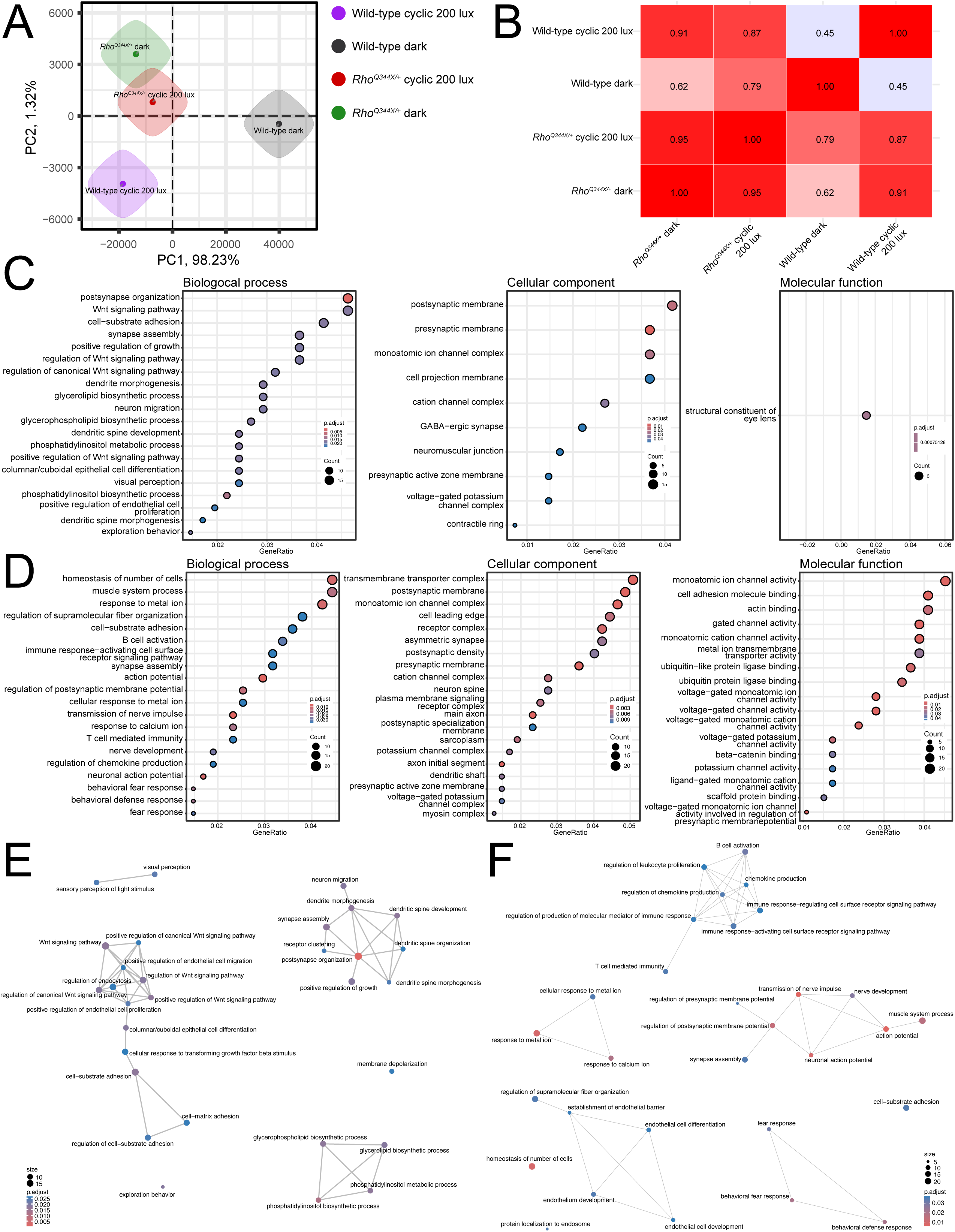
RNA-seq analysis of P35 wild-type and *Rho^Q344X/+^* mouse retinas indicates that light exposure has minimal impact on the transcriptome of *Rho^Q344X/+^* retinas. (A) Principal component analysis (PCA) shows that wild-type retinas reared under standard cyclic 200 lux light (purple) and constant dark (gray) conditions cluster separately. *Rho^Q344X/+^* retinas reared under standard cyclic 200 lux light (red) and constant dark (green) conditions cluster separately from either of their wild-type counterparts. *Rho^Q344X/+^* retinas cluster similarly between standard cyclic and constant dark conditions. (B) The table presents the Pearson correlation coefficients among the experimental groups, as indicated along the left and bottom margins. (C and D) Pathway analyses for gene ontology (GO) terms using mRNA expression data filtered for a 2-fold change were performed for wild-type and *Rho^Q344X/+^* mice (C and D, respectively) reared under standard cyclic 200 lux lighting or darkness. Significantly altered pathways (up to 20) are shown for biological process (left most), cellular component (middle), and molecular function (right most). The size of each dot represents the level of protein enrichment (Count), and the color coding indicates the statistical significance (p.adjust), as indicated on the right side of the panels. (E and F) Networks illustrating the output of the hypergeometric tests. The pathway analyses were conducted using the gene enrichment data for wild-type (E) and *Rho^Q344X/+^* mice (F) under two different conditions, standard cyclic 200 lux lighting and darkness. The size of each dot (Count) represents the level of gene enrichment (number of genes in each pathway), and the color coding shows statistical significance (p.adjust), as indicated in the bottom left (E) and bottom right (F) corners of the panels.

To understand the specifics of light-dependent changes, we conducted pathway analysis employing GO terms. In dark-reared wild-type mice (Supplemental Figure S4), genes involved in visual perception such as sensory perception of light stimuli (GO0050953) and visual perception (GO0007601) were upregulated compared to wild-type mice reared under standard cyclic 200 lux light condition (Figure 3C and E). Those are mainly crystallins, such as Cryaa (crystallin, alpha A, 2395% 200 lux light), Cryba1 (crystallin, beta A1, 2217.1% 200 lux light), and Crybb3 (crystallin, beta B3, 6939.3% 200 lux light), that function as structural constituent of eye lens (GO0005212) (Figure 3C, right-most panel). They also function as chaperone molecules and are known to be regulated by light or circadian rhythm in rats (21, 22). In addition, components involved in synapse maturation (e.g., GO0099173) were downregulated, whereas components involved in WNT signaling (e.g., GO0030177) were either upregulated or downregulated. The downregulation in the synapse maturation pathway is consistent with the previous observation that synapse maturation is delayed under a dark condition (23). Upregulation of molecules involved in the WNT signaling such as Wnt5a (wingless-type MMTV integration site family, member 5A, 252.8% 200 lux light), Fgf2 (fibroblast growth factor 2, 1602% 200 lux light), and Cav1 (caveolin 1, caveolae protein, 240.4% 200 lux light) are consistent with light-dependent regulation of developing retinal architectures, including its vasculatures (24, 25).

In the *Rho^Q344X/+^* mice, through pathway analysis employing GO terms (Supplemental Figure S5), genes involved in inflammation such as homeostasis of number of cells (GO0048872) and B cell activation (GO0042113) changed their expression levels (Figure 3D and F). In addition, pathways involved in synapse maturation such as synapse assembly (GO0007416) and regulation of postsynaptic membrane potential (GO0060078) were upregulated in the dark-reared *Rho^Q344X/+^* mice.

For light-dark differences, more GO terms were identified in *Rho^Q344X/+^* mice than in wild-type mice (Figure 3C and D), although the PCA and Pearson correlation analyses indicated less pronounced transcriptomics changes in *Rho^Q344X/+^* mice than in wild-type mice (Figure 3A and B). This apparent discrepancy can be attributed to the overrepresentation of GO terms associated with inflammatory pathways, whose expression changes were predominantly observed in *Rho^Q344X/+^* mice.

While these physiological light conditions did not exacerbate the degree of photoreceptor degeneration, we found slight exacerbation of photoreceptor degeneration in *Rho^Q344X/+^* mice when we employed more severe light stimulation, at an intensity of 3000 lux for a continuous 24-hour light cycle over 5 days (constant 3000 lux), as observed by OCT (Figure 4A and B). The 2500 lux cyclic light conditions we introduced in this study did not cause degeneration of rod photoreceptors in wild-type mice as described in previous studies (18–20). The effect of constant 3000 lux light on *Rho^Q344X/+^* mice was variable among the four quadrants of the retina. In the nasal and temporal retinas, ONL was thinner under constant 3000 lux light condition (82.3 – 89.4%) than under other light conditions (Figure 4B). Intriguingly, we did not observe significant retinal thinning in the ventral and dorsal regions due to constant 3000 lux light exposure (Figure 4B). Detailed analyses of histology confirmed the absence of light effects in worsening photoreceptor degeneration in the ventral region (Figure 5A and B, dark vs constant 3000 lux). In *Rho^Q344X/+^* mice reared under constant 3000 lux light condition, slight thinning or thickening of ONL was observed in a few dorsal regions, however, no significant changes in ONL thicknesses were observed in the ventral region. Likewise, OS layer thicknesses were similar among these different light conditions (Figure 5B, right panel). Under the same conditions, wild-type mice did not demonstrate significant thinning of the ONL as a result of 3000 lux light exposure (Figure 5C and D, dark vs constant 3000 lux). Intriguingly, the OS layer of 2500-lux-light-exposed wild-type mice showed significant thinning in the ventral region and a few dorsal regions (Figure 5D, right panel), consistent with the mRNA downregulation of the OS component, *Rho*, observed by qPCR (Figure 1F, blue bars). Taken together, these results indicate that light does not significantly affect retinal degeneration in *Rho^Q344X/+^* mice.

**Figure 4.**
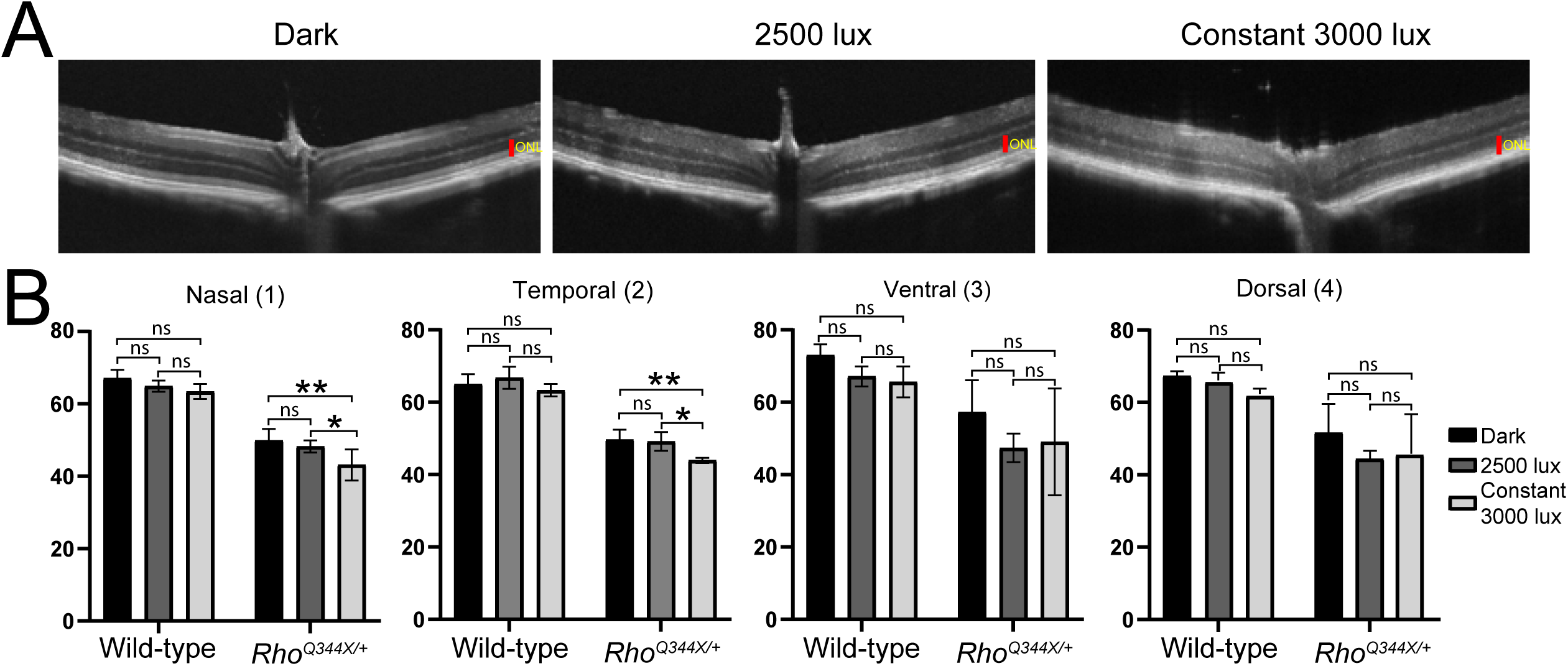
OCT shows photoreceptor degeneration in the *Rho^Q344X/+^* knock-in mice under constant 3000 lux light condition. (A) OCT images spanning the ventral-ONH-dorsal regions were acquired for P35 *Rho^Q344X/+^* mice exposed to constant 3000 lux light for 5 consecutive days. ONLs are indicated by red vertical bars. Scale bars, 100 µm. (B) ONL thicknesses were measured for the OCT images from P35 wild-type or *Rho^Q344X/+^* mice subjected to 3 different light conditions, dark-reared (black), cyclic-2500 lux-reared (dark gray), and constant 3000 lux (light gray). The data are presented as mean ± SD (n = 4 animals for each genotype). The data were subjected to statistical analyses using two-way ANOVA, followed by Tukey’s post-hoc test for pairwise comparisons. Statistical significance (**, *p* < 0.005; *, *p* < 0.01; ns, not significant) is indicated above each pair.

**Figure 5.**
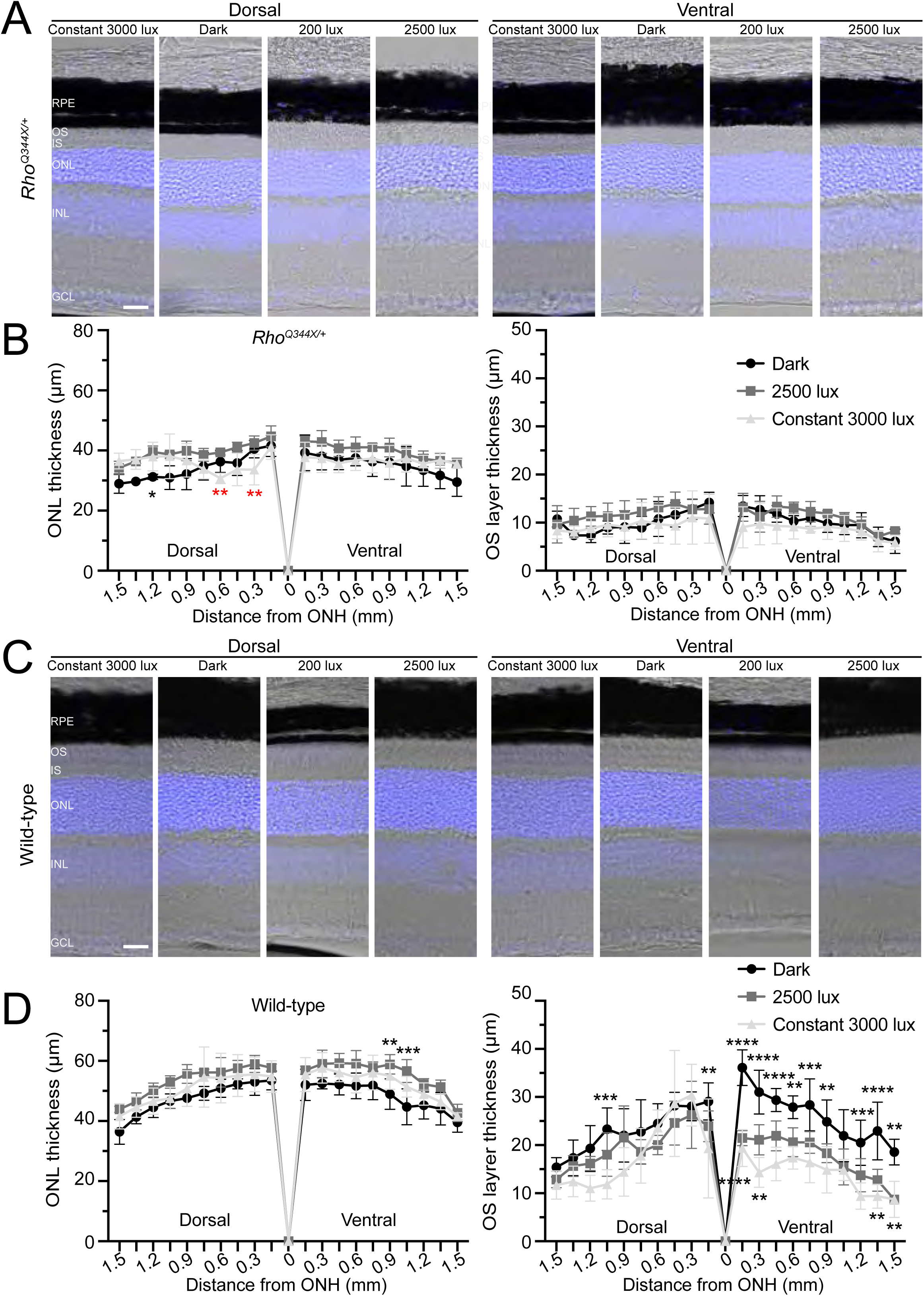
OS layer thickness and degrees of rod photoreceptor degeneration are similar in *Rho^Q344X/+^* mice regardless of light conditions. (A and C) Retinal sections from P35 *Rho^Q344X/+^*(A) and wild-type (C) mice reared under one of the following conditions were labeled with Hoechst 33342 (blue) to visualize their nuclear layers: 5 day constant exposure to 3000 lux, dark, standard 200 lux 12L/12D cyclic light condition, or 2500 lux cyclic 12L/12D light condition (from left to right). Dorsal and ventral retinas were subjected to imaging (left and right panels, respectively). (B and D) Thicknesses of the ONL and OS layers were measured every 150 µm on the dorsal and ventral sides of the ONHs for *Rho^Q344X/+^* (B) and wild-type (D) mice reared under 5-day constant exposure to 3000 lux (light gray triangle), dark (black circle), and 2500 lux cyclic 12L/12D light conditions (dark gray rectangle). The data show ONL and OS layer thicknesses differ in some regions of wild-type and *Rho^Q344X/+^* retinas. The data were analyzed using two-way ANOVA and are shown as mean ± SD (n = 4 for each light condition). For comparisons among light conditions, Tukey’s post hoc test for pairwise comparisons was used to calculate p-values. Statistical significance (****, *p* < 0.0001; ***, *p* < 0.001; **, *p* < 0.005; *, *p* < 0.01) is indicated above or below each data point between dark and 2500 lux (above each data point), between dark and constant 3000 lux (below each data point), and between cyclic 2500 lux and constant 3000 lux (in red below each data point).

### Photoreceptor functions are attenuated in *Rho^Q344X/+^* mice

In humans, the *Rho^Q344X^* mutation causes progressive visual impairments in an autosomal dominant manner (26). To study the nature of photoreceptor dysfunction, we conducted electroretinogram (ERG) analyses of *Rho^Q344X/+^* mice, using wild-type mice as controls (Figure 6). In *Rho^Q344X/+^* mice at P21, we found that a-wave responses, reflective of rod function, were significantly lower at light intensities ranging from -0.7 to 1.6 log(cd·s/m²) (Figure 6A and B). From P21 to P35, the a-wave amplitudes declined significantly (Figure 6A - C). The amplitudes of *Rho^Q344X/+^* mice were 41.6, 55.3, 52.5% (41.6-55.3%) of those in wild-type mice at the light intensities of -0.7, 0.3, and 1.6 log(cd·s/m^2^) at P21 (Figure 6B, top graph). The responses of *Rho^Q344X/+^*mice were 34.4, 41.5, and 40.4% (34.4-41.5%) of those in wild-type mice at the light intensities of -0.7, 0.3, 1.6 log(cd·s/m^2^) at P35 (Figure 6C, top graph). B-wave responses were less affected during this time course (Figure 6B and C, bottom graphs). While b-wave amplitudes showed a lower trend in *Rho^Q344X/+^* compared to wild-type mice, those differences were not statistically significant (*p* > 0.1).

**Figure 6.**
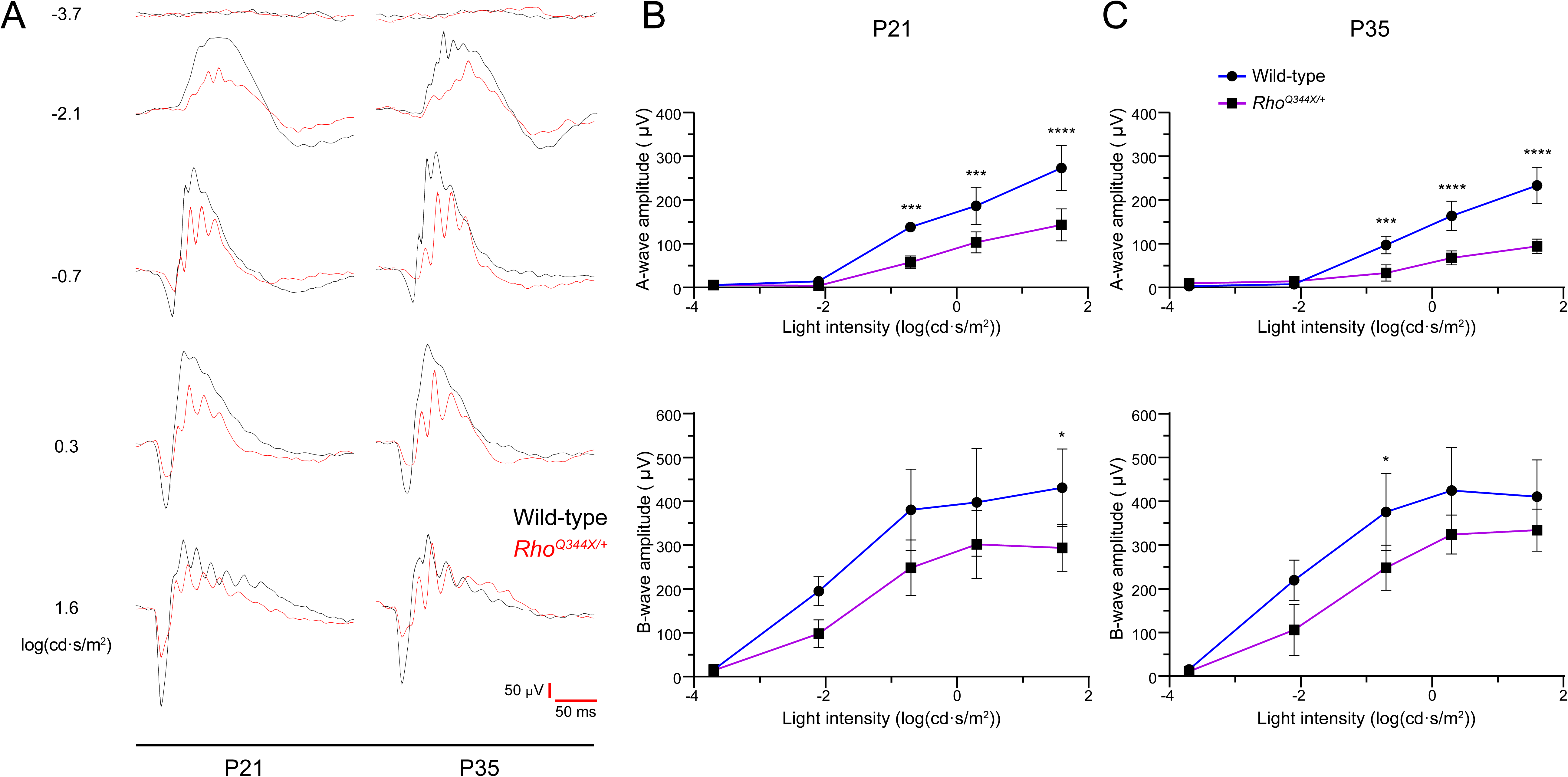
Electroretinogram (ERG) shows that light response is impaired in the class I *Rho^Q344X^* knock-in mutant mouse model. (A) Representative scotopic ERG responses were measured at 5 different light intensities represented by log(cd·s/m^2^). The responses were recorded from wild-type (black line) and *Rho^Q344X/+^* (red line) mice reared under standard cyclic 200 lux conditions and aged at P21 or P35. Response time (millisecond, ms) and amplitude (μV) are indicated by red bars at the bottom right corner of the panel. (B and C) Amplitudes of scotopic a- and b-waves from P21 (B) or P35 (C) wild-type (blue) and *Rho^Q344X/+^* (purple) mice were plotted as a function of light intensities (top and bottom, respectively). The data are represented as mean ± SD (n = 4 animals for each genotype) and were subjected to statistical analysis using two-way ANOVA, followed by Šídák’s multiple comparisons test for pairwise comparisons. Statistical significance (****, *p* < 0.0001; ***, *p* < 0.001; \**p* < 0.05) is indicated above each data point.

We investigated whether rearing *Rho^Q344X/+^* mice under high-intensity light conditions of cyclic 2500 lux accelerates any photoreceptor dysfunction, thereby decreasing electrophysiological responses detectable by ERG (Figure 7). We observed similar a- and b-wave forms (Figure 7A) and their amplitudes between dark-reared and 2500-lux-light-reared animals (Figure 7B, *p* > 0.5, not significant). These results were consistent with the lack of structural change due to 2500 lux light exposure, indicating that ectopic rhodopsin activation under physiological light intensity does not cause degeneration or physiological dysfunction of rod photoreceptors.

**Figure 7.**
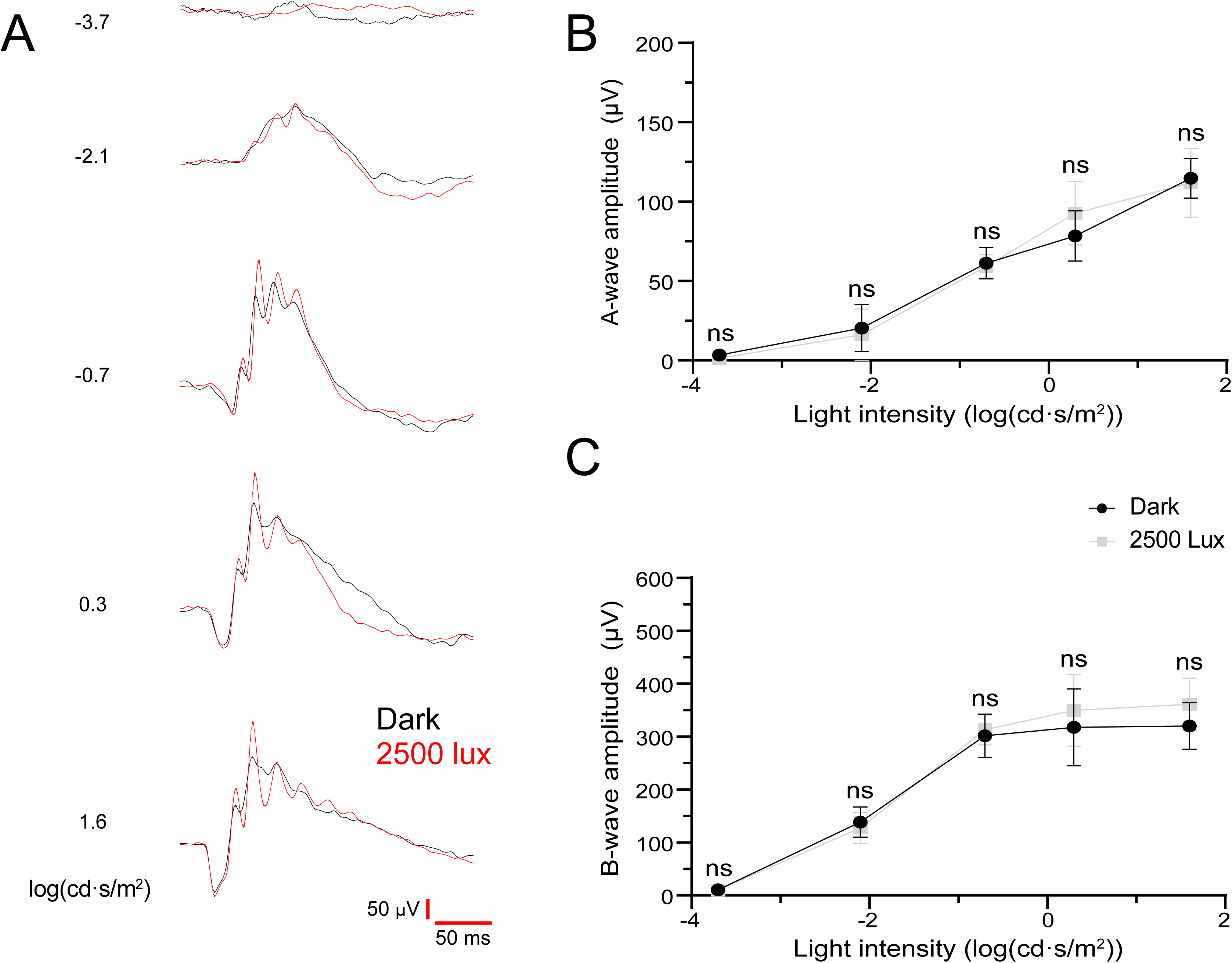
Scotopic ERG responses from P35 *Rho^Q344X/+^* mice are similar under both dark and cyclic 2500 lux rearing conditions. (A) Representative scotopic ERG responses were measured at 5 different light intensities represented by log(cd·s/m^2^). The responses were recorded from P35 *Rho^Q344X/+^* mice reared under dark (black line) or 2500 lux cyclic lighting (red line) conditions. Response time (millisecond, ms) and amplitude (μV) are indicated by red bars at the bottom right corner of the panel. (B) Scotopic a- and b-wave amplitudes were measured from P35 *Rho^Q344X/+^* mice reared under darkness (black circle) or cyclic 2500 lux (gray square), and plotted as a function of light intensities (top and bottom, respectively). The data are presented as mean ± SD (n = 4 animals for each genotype) and were subjected to statistical analysis using two-way ANOVA, followed by Šídák’s multiple comparisons test for pairwise comparisons. Neither a- nor b-wave amplitudes showed statistical significance. ns, not significant.

### Proteins involved in secretory pathways are downregulated in *Rho^Q344**X**/+^* retinas

To understand complex global biological changes in the *Rho^Q344X/+^*retina, we quantitatively compared mRNA and protein levels using RNA-seq and our previous mass spectrometry data (Figure 8) (4), respectively. Pearson correlation between transcriptomic and proteomic data in P35 wild-type retinas was 0.495, similar to the value observed for protein-mRNA expressions in previous studies (27). Pearson correlation between transcriptomic and proteomic data in P35 *Rho^Q344X/+^*retinas was 0.500, which was a slightly better correlation value than that of wild-type retinas. Those analyses indicate that, in general, retinal mRNA and protein expression levels demonstrate a modest degree of correlation.

**Figure 8.**
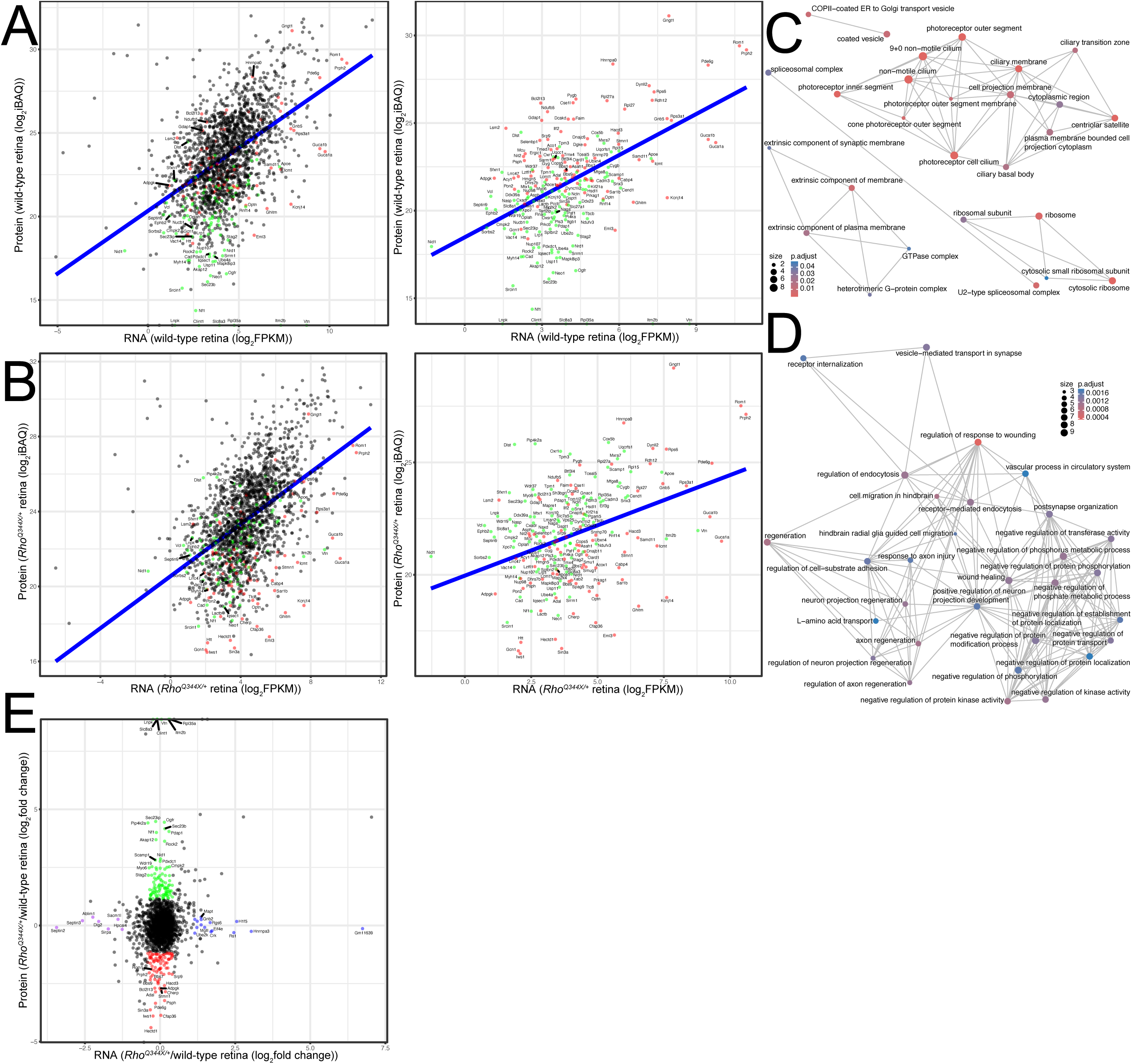
Comparison between proteomics and RNA-seq for P35 wild-type and *Rho^Q344X/+^* retinas shows distinct downregulation and upregulation. (A and B) Protein expression levels were plotted against RNA expression levels. Among the gene products that remained unchanged in the RNA-seq data (± 1.33-fold or 75-133.3% of control), proteins that were significantly downregulated (< 45%) or upregulated (> 222%) in the proteomics analysis (indicating over a 2.22-fold change) are indicated as red (downregulated) and green (upregulated) dots, respectively. Proteins whose expression levels did not change significantly are shown as black dots. Plots are for wild-type (A) and *Rho^Q344X/+^* (B) retinas. The right panels display the data with the black dots omitted. A blue line in each panel indicates the linear regression. (C and D) Networks illustrating the outcomes of the hypergeometric test. The pathway analysis was based on gene enrichment for gene ontology terms. Networks for downregulated proteins (represented in red in (A) and (B)) are presented in (C), whereas networks for upregulated proteins (represented in green in (A) and (B)) are presented in (D). The size of each dot (Count) represents the level of gene enrichment (number of genes in each category), while the color coding indicates statistical significance (p.adjust), as indicated at the bottom left (C) and top right (D) corners of the panels. (E) Log2-transformed fold changes from proteomic data were plotted against log2-transformed fold changes from RNA-seq data. Proteins that showed differential expression only in proteomics data are represented by red dots (for downregulated proteins) and green dots (for upregulated proteins). Gene products that show differential expression only in RNA-seq are represented by purple dots (for downregulated mRNA) and blue dots (for upregulated mRNA), respectively. Proteins whose expression levels did not change significantly are shown as black dots.

Our studies indicated a dramatic reduction in rhodopsin transport of *Rho^Q344X/+^* rod cells. Rhodopsin is the most highly expressed cargo in the endoplasmic reticulum (ER)-Golgi and post-Golgi secretory pathways of wild-type rods. We hypothesized that a major disruption in the flow of these pathways would have a significant impact on global protein homeostasis, particularly on proteins that are involved in the secretory events, while exerting minimal effects on mRNA levels. To test this hypothesis in an unbiased manner, we implemented stringent cutoff criteria to define mRNA transcripts that did not change their expression levels significantly. Specifically, mRNA transcripts with expression levels within ± 1.33-fold (75-133%) of wild-type levels were considered unchanged. To define proteins with significant expression changes, we implemented more stringent cutoff criteria: a 2.22-fold change (< 45% or > 222%) relative to wild-type levels was considered significantly downregulated or upregulated. Based on these criteria, 1693 proteins showed no significant alterations in abundance when their corresponding mRNA transcripts also remained unchanged (Figure 8A, B, and E, black dots). This analysis supports that the expression levels of most proteins are correlated with their respective mRNA levels. Among proteins without significant mRNA changes, 85 were downregulated (Figure 8A, B, and E, red dots), whereas 101 were upregulated (Figure 8A, B, and E, green dots). Those proteins were subjected to GO term pathway analyses to understand which pathway components were either downregulated or upregulated. Among the downregulated GO terms were photoreceptor OS and cilia (e.g., GO0097733 and GO0060170), GTPase complex (e.g., GO1905360), spliceosomal complex (e.g., GO0005681), ribosome (e.g., GO0005840), and COPII-coated ER vesicle (e.g., GO0030134) (Figure 8C). More specifically, in photoreceptor OS and cilia pathways, following proteins were significantly downregulated: G_β_5 (guanine nucleotide binding protein (G protein beta 5, or Gnb5) (43.6% wild-type), rod transducin γ subunit (G protein gamma transducing activity polypeptide 1, or Gngt1) (26.5% wild-type) (5), rod outer segment membrane protein 1 (Rom1, 27.0% wild-type), Prph2 (24.4% wild-type), and Bardet-Biedl syndrome 7 (Bbs7, 22.8% wild-type) proteins. While these proteomics results were consistent with our previous study (4), these new analyses indicate that rhodopsin is required for the synthesis or stability of other rod OS proteins. In the COPII-coated ER vesicle pathway, the following proteins were downregulated: endoplasmic reticulum-golgi intermediate compartment 1 (Ergic1, 37.7% wild-type) involved in the turnover of Ergic2 and Ergic3 (28), secretion associated Ras related GTPase 1b (Sar1b, 40.2% wild-type) involved in the COPII coat assembly (29–31), and transmembrane p24 trafficking protein 7 (Tmed7, 24.4% wild-type) known to be a COPII adapter mediating vesicular protein transport (32, 33).

In the ribosome GO term, Abce1 (ATP-binding cassette, sub-family E member 1, 42.4% wild-type) associated with the initiation of translation and ribosomal recycling (34), and Gcn1 (GCN1 activator of EIF2AK4, 20% wild-type) associated with translational quality control and mRNA decay (35, 36), were downregulated. Related to Gcn1 downregulation, we found that its interactor, Rnf14 (ring finger protein 14, 248.3% wild-type), known to ubiquitinate Gcn1 (37), was upregulated. Upregulation of Rnf14 results in increasing degradation of eukaryotic translation elongation factor 1A and affects translation (38, 39). These downregulated pathways may partially account for the alterations in protein expression levels despite the absence of changes in mRNA expression.

Among the upregulated GO terms were axon neuron regeneration projection (e.g., GO0042060), receptor-mediated endocytosis (e.g., GO0006898), vascular process in circulatory system (e.g., GO0003018), negative protein kinase activity (e.g., GO0042326), and L-amino acid transport (e.g., GO0015807) (Figure 8D). In these pathways, the following proteins were upregulated without significant changes in their mRNA expressions: Rock2 (Rho-associated coiled-coil containing protein kinase 2, 1236.6% wild-type) involved in microglial proliferation (40), and Nf1 (neurofibromin 1, 1606% wild-type) known to modulate microglial function (41). In addition to these proteins, Itm2b (integral membrane protein 2B) known to be highly expressed in microglia (42) was only detected in *Rho^Q344X/+^*mice. Also, expression levels of the following proteins increased remarkably: Pip4k2a (phosphatidylinositol-5-phosphate 4-kinase, type II, alpha, 2136% wild-type) known to be expressed in microglia/macrophage (43) and regulate intracellular cholesterol transport (44), Sec23ip (Sec23 interacting protein, 2232% wild-type) known to be involved in cholesterol trafficking (45), and Ogfr (opioid growth factor receptor, 2180% wild-type) known to be involved in cytokine production (46, 47) and lipid oxidation (48). These observations indicate that microglial activation involves translational or posttranslational mechanisms to upregulate proteins (49, 50).

## Discussion

In *Rho^Q344X/+^* mice, mutant and wild-type rhodopsin are both significantly downregulated prior to the onset of rod degeneration. This downregulation occurs at the level of mRNA. The degree of *Rho* mRNA downregulation (46.0% of wild-type) is consistent with previous immunoblotting results, which showed significant downregulation of rhodopsin (37.5% of wild-type) in *Rho^Q344X/+^* mice (4). To a certain degree, we believe that the previously described lysosome-mediated degradation of Rho^Q344X^ proteins contributes to the loss of rhodopsin at the protein level (4, 51). Nevertheless, given that rhodopsin protein levels and gene dosage are linearly correlated (2), the evidence strongly suggests that the dramatic mRNA downregulation is the primary contributor to rhodopsin downregulation in *Rho^Q344X/+^*mice. Considering knock-in adRP models, as introduced in this study, express wild-type and mutant mRNAs at equivalent levels (3, 4), the dramatic downregulation attenuates the normal function of rhodopsin as well as the toxicities of mutant rhodopsin molecules. In *Drosophila melanogaster*, an adRP causative rhodopsin mutation not only facilitates the degradation of the mutated proteins but also promotes the degradation of the wild-type protein, significantly promoting photoreceptor degeneration (52). The existence of such dominant negative effects at the protein level is likely a minor contributor in mammals. Instead in mice, we discovered a novel dominant-negative mechanism that acts at the mRNA to impact the quantity of rhodopsin. As we observe similar rhodopsin downregulation in another rhodopsin-RP model, *Rho^P23H/+^* mice, the dominant-negative effect appears to be common among class I and class II rhodopsin-RP models.

This study indicates that light does not exacerbate rod degeneration in *Rho^Q344X/+^* mice. Ectopic rhodopsin activation has been demonstrated in previous studies using animal or cell culture models of rhodopsin mislocalization (13, 15, 16). In our study, the light had a minimal impact on rod photoreceptor degeneration of *Rho^Q344X/+^* mice at established cyclic light intensities of 200–2500 lux which are typical for daily activities such as office work. Furthermore, light at 200 lux had a minimal impact on the retinal transcriptome, in contrast to the significant effects observed in wild-type mice. The significant transcriptomic changes observed in wild-type mice are consistent with the known effects of light on retinal developmental processes, including those of synapses and vasculature (23–25). Previous studies of transgenic mice expressing *Rho^Q344X^* demonstrated a marked exacerbation of rod degeneration under continuous exposure to 3,000 lux for five days (13). Therefore, we tested a similar extent of light exposure on our *Rho^Q344X/+^* knock-in mice and found that it minimally affected the retinal architecture. Mutant rhodopsin is known to cause rod degeneration in a dosage-dependent manner (8, 53, 54). The previous transgenic *Rho^Q344X^* mice demonstrate rod degeneration that is more rapid than that observed in our *Rho^Q344X^*knock-in mice, signifying higher expression of *Rho^Q344X^* in individual rods of the prior models (4, 13). Given that rhodopsin serves as the chromophore involved in light-dependent rod degeneration (13, 15, 16, 19), the dramatically attenuated expression of *Rho* mRNA—along with the corresponding decrease in rhodopsin protein levels (4)—is likely a key contributor to the apparent lack of light response observed in our study, as evidenced by ERG measurements.

The extent to which light impacts the survival of rod photoreceptors across a broader spectrum of rhodopsin-RP remains unclear. Unlike our observations for class I rhodopsin mutants, other classes of rhodopsin mutations are thought to trigger light-dependent toxicity, which exacerbates photoreceptor degeneration and leads to severe conditions such as sector RP in humans, as well as similar localized degenerative conditions in animal models (55, 56). The P23H mutation, which causes rhodopsin misfolding, and the T4R and N15S mutations, which lead to defects in rhodopsin glycosylation, are particularly significant in this context. The T4R mutation in a canine model results in sector RP, while the P23H knock-in mouse model exhibits more severe rod degeneration in the ventral retina compared to the dorsal region (11, 57). As areas of degeneration coincide with increased light exposure, it is believed that this occurs due to the light activation of mutant rhodopsin, resulting in its misfolding or aberrant activation (58, 59). Unlike these mutations affecting folding and glycosylation, there are no reports associating rhodopsin class I mutations with sector RP, to the best of our knowledge. Addressing the role of light as an environmental risk factor is critical when considering mutation-specific management strategies for RP patients, particularly in determining safe levels of light tolerance that can be endured without exacerbating irreversible blindness. In the case of class I mutation, basal downregulation of *Rho* mRNA and further downregulation by light are the contributing factors attenuating light-dependent toxicity.

While rhodopsin downregulation is mainly due to its mRNA downregulation, various proteins were downregulated without significant changes in their mRNA levels. Those downregulated proteins are associated with COPII-coated ER vesicle, photoreceptor OS and cilia, among others. Downregulation of components associated with COPII-coated ER vesicle pathway included Sar1, which is associated with ER to Golgi transport of rhodopsin (60). It is likely that decreased flow of rhodopsin in this pathway led to downregulation of these components as part of adaptation. Sar1 is involved in the ciliary transport of Peripherin-2 (61), which is downregulated at the protein level without significant changes in its mRNA levels. Thus, reduced flow of ER-to-Golgi and Golgi-to-cilia transport partly explains why some of the OS proteins are downregulated at the protein levels. Likewise, regarding Golgi-to-ciliary transport, BBS components are downregulated at the protein levels. Tmed7, a COPII adapter protein, is involved in the ER-Golgi transport of specific cargoes in the secretory pathway (33). *TMED7* homozygous mutation is associated with inherited retinal degeneration (62), suggesting its involvement in the secretory pathway of retinal cells. A decline in rhodopsin flow, due to transcriptional downregulation, eventually propagates at the protein levels to downregulate various components of vesicular transport machinery at the translational or posttranslational levels.

In summary, we identified a novel transcript-level dominant-negative effect exerted by mutant *Rho* alleles, which significantly influences the expression of wild-type rhodopsin. This mechanism introduces a previously uncharacterized layer of complexity in the pathology of rhodopsin-RP. Although our primary focus was a class I mutant, we observed that a prominent class II mutant exhibits a similar mRNA-level effect (this study) and corresponding protein-level rhodopsin downregulation (63), suggesting that the dominant negative effect is a shared outcome in RP. This finding aligns with the hallmark RP phenotype characterized by the shortening and loss of rod OSs (64). These observations raise the possibility that rhodopsin downregulation may represent an early and common step in the progression of vision loss in RP patients. Our results also contrast with the previously described protein-level dominant-negative effect observed in *Drosophila*, where mutant rhodopsin proteins destabilize wild-type proteins via the unfolded protein response and autophagy of ER (52). Instead, our findings highlight mRNA regulation that has dual implications: these mechanisms might mitigate the toxicity of the mutant allele while simultaneously compromising the function of the wild-type allele. Further investigation into the mechanism underlying *Rho* mRNA regulation is warranted. For adRP caused by rhodopsin mutations (rhodopsin-RP), combined suppression and replacement gene therapies have been proposed. This approach involves RNA interference to suppress endogenous rhodopsin genes, including the mutant allele, while introducing a wild-type rhodopsin gene resistant to suppression (65). Multiple lines of evidence indicate that rhodopsin downregulation is an effective strategy to mitigate rod degeneration associated with other gene mutations (66, 67). Our findings suggest that rod photoreceptors possess endogenous mechanisms for such suppression, potentially mirroring aspects of RNAi-based therapy. Exploring these natural regulatory mechanisms could inform the development and refinement of gene therapies, paving the way for more effective treatment strategies for RP.

## Methods

### Animals

All animal experiments conducted adhered to the procedures approved by the Institutional Animal Care and Use Committee (IACUC) at Indiana University School of Medicine and were in compliance with the guidelines set forth by both the American Veterinary Medical Association Panel on Euthanasia and the Association for Research in Vision and Ophthalmology. Wild-type and *Rho^Q344X^* mutant mice on the pigmented C57BL/6 background were housed under a 12-hour light/12-hour dark (7 a.m./7 p.m.) cycle and received standard mouse chow. For constant dark or 2500 lux 12-hour light/12-hour dark (7 a.m./7 p.m.) light cycle, mice were reared in the circadian cabinet (Actimetrics, Wilmette, IL, USA) from P0 until the time points of experiments. For exposure to 5 day constant 3000 lux light condition, we essentially followed the described protocol in the previous study (13). In brief, pigmented wild-type and *Rho^Q344X/+^* mice were housed under a 12-hour light/12-hour dark (7 a.m./7 p.m.) cycle and received standard mouse chow after birth until P29. On P29, mice were exposed to constant 3000 lux light intensity (24 hours) for 5 days. Then mice were housed one more day under a 12-hour light/12-hour dark with standard ambient light (200 lux) condition before the OCT and collecting the eyes for sections. For the housing light intensities, light intensities were adjusted to the experimental light conditions using light meter. We confirmed that the mice used in this study do not carry rd1 and rd8 mutations (68, 69).

### Estimation of retinal protein abundance

To evaluate the relationship between transcript and protein levels in the retina, we estimated retinal protein abundance using iBAQ (intensity-based absolute quantification) values calculated with MaxQuant (70). These values were derived from our previously published dataset, available through the ProteomeXchange Consortium via the MassIVE partner repository (dataset identifier PXD046795) (4). For the combined analyses of transcriptomic and proteomic data, a 2.22-fold change (< 45% or > 222%) relative to wild-type levels was considered indicative of significant protein downregulation or upregulation, while changes within ± 1.33-fold (75-133%) of wild-type mRNA levels were considered insignificant. Pathway analyses were performed in R (version 3.3.0+) using the clusterProfiler package (30) via the RStudio interface (version 4.3.1). GO terms with corrected P values of less than 0.05 were considered significantly enriched with differentially expressed proteins.

### Transcriptomics analysis using RNA-seq

Total RNA was extracted and pooled from mouse retinas from P35 wild-type and *Rho^Q344X/+^* mice reared under standard cyclic 200 lux light or dark conditions (n = 4 animals for each group). RNA quality was checked by Bioanalyzer (Agilent Technologies) and spectrophotometer. Using these total RNA samples, RNA-seq libraries were prepared with the NEBNext Ultra II Directional RNA Library Prep kit for Illumina (New England Biolabs). RNA-seq was contracted to LC Sciences (Texas, USA). The RNA-seq libraries were subjected to sequencing by NovaSeq6000 (Illumina Inc., San Diego, CA, USA). Obtained sequence data, collections of 150 nt paired-end short reads, were mapped to the C57BL mouse genome reference sequence database (GRCm39) with HiSAT2 (71). GO enrichment analysis of differentially expressed genes was implemented by GOseq (72), through which gene length bias was corrected. GO terms with corrected P values of less than 0.05 was considered significantly enriched with differentially expressed genes. PCA analysis, Pearson correlation, and pathway analyses were performed in R (version 3.3.0+) using the PCAtools, Stats, and clusterProfiler package (30), respectively, via the RStudio interface (version 4.3.1).

### Examination of *Rho* mRNA expression levels in the wild-type, *Rho^Q344X^*and *Rho^P23H^* mutant retinas

Total retinal RNA was extracted from wild-type, *Rho^Q344X^*, and *Rho^P23H^* mutant mice at three different time points (P14, P21, and P35). For each sample, two µg of total RNA was reverse transcribed into cDNA using SuperScript IV reverse transcriptase (Thermo Fisher Scientific). Ten ng of resultant cDNA served as a template for qPCR. This qPCR assay employed a SYBR Green-based protocol (73) using QuantStudio™ 6 Flex Real-Time PCR System (Thermo Fisher Scientific) following the manufacturer’s protocol. The following primer pair was used for *Rho*: forward primer, 5’-CTGTAATCTCGAGGGCTTCTTT-3’; reverse primer, 5’-GTGAAGACCACACCCATGATAG-3’.

The following primer pair was used for *Gnat1*: forward primer, 5’-GGACGACGAAGTGAACCGAATG-3’; reverse primer, 5’-TAGTTGCCGGCATCCTCGTAAG-3’.

The following primer pair was used for *Pde6a*: forward primer, 5’-CCCAACACAGAAGAGGATGAGC-3’; reverse primer, 5’-TCTCTTGGTGAAGTGGGGTTCA-3’.

The following primer pair was used for actin (*Actb*): forward primer, 5’-GAACATGGCATTGTTACCAACT-3’; reverse primer, 5’-TCAAACATGATCTGGGTCATCT-3’.

The temporal expression levels of cDNAs were plotted for wild-type, heterozygous, and homozygous mutant mice at P21, P28, and P35.

### OCT

OCT images were acquired as described (4). In brief, wild-type or *Rho^Q344X^* mutant mice were anesthetized with isoflurane, and their pupils were dilated using 0.5% tropicamide and 2.5% phenylephrine. OCT images of these mice were captured at P21, P35, P60, and P120, using the Phoenix Research Labs Reveal Optical Coherence Tomography (OCT2) Imaging System (Micron IV, Phoenix Research Laboratories, Pleasanton, CA, USA). To evaluate the total retinal thickness and ONL thickness, cross-sectional retinal images passing through the optic nerve head (ONH) were obtained in the dorsal-ventral and nasal-temporal axes. OCT images were captured using the line type and full line size settings with an average of 60 frames/scan. Enhanced Depth Imaging was selected to improve the signal originating from the outer retina. The acquired OCT images were segmented for retinal layers and analyzed using InSight software (Voxeleron LLC). The nasal, temporal, ventral, and dorsal regions, located 500 µm from the ONH, were used for the comparison.

### Morphometry by light microscopy

For histological analysis of the retina, we followed the protocol described in the previous study (4). In brief, wild-type and *Rho^Q344X/+^*mice were sacrificed at P35, and their eyes were enucleated. Eyecups were prepared and fixed in 4% paraformaldehyde/PB for 6 hours, washed by PBS 3 times, and embedded in 1.5% agarose. Eyecup sections with a thickness of 50 µm were prepared using a vibratome (7000 SMZ, Campden Instruments, UK). Sections were cut in the dorsal-ventral axis, and those passing through the ONH were used for further analysis. The samples were stained with Hoechst 33342 (50 ng/mL in PBS) and mounted on glass bottom dish with VECTASHIELD® PLUS Antifade Mounting Medium (Vector Laboratories). Images were acquired with AX R confocal microscope (Nikon) using 20x Apo Lambda S (NA = 0.95) water immersion objective lens. The thicknesses of the ONL were measured in four sections prepared from four independent mice for each genotype and time point. Thicknesses of the ONL were measured at 150 µm intervals using NIS-Elements (Nikon).

### ERG

As described above, *Rho^Q344X/+^* mice were reared in the dark, under cyclic 200 lux lighting, or under cyclic 2500 lux lighting (n = 4 animals for each group). Wild-type mice were reared under cyclic 2500 lux lighting (n = 4 animals for each group). After rearing mice under these conditions, they were dark-adapted overnight and were subjected to ERG at either P21 or P35. Wild-type or *Rho^Q344X/+^* mice were anesthetized with 90 mg/kg of ketamine and 10 mg/kg xylazine, and their pupils were dilated using 0.5% tropicamide and 2.5% phenylephrine. Mice were placed on the stage with a heat pad (LKC Technologies, Gaithersburg, Maryland, USA). The heat pad was set to 37°C. Scotopic ERG responses were recorded for 5 light intensities (−3.7, -2.1, -0.7, 0.3, 1.6 log(cd·s/m^2^). Amplitudes of a- and b-waves were manually determined using EMWIN software (LKC Technologies).

## Statistical analysis

Statistical analyses and data visualization, except for the proteomics and transcriptomics data, were conducted using GraphPad Prism software (version 8.0). Unless otherwise specified in the figure legends, one-way ANOVA with Tukey’s post hoc test was used for multi-group comparisons, and an unpaired two-tailed t-test was used for two-group comparisons. *P*-values less than 0.05 were considered significant. The data are presented as the mean ± SD.

## Acknowledgements

We thank Ms. K. C. Lynn and Ms. K. A. Hansen for their technical assistance in the maintenance and genotyping of the mice used in this study. We thank Dr. S. Imanishi for her aid and assistance to the project. Mass spectrometry data were acquired by the Proteomics and Metabolomics Core at the Lerner Research Institute of the Cleveland Clinic Foundation. We thank Drs. Belinda Willard and Ling Li for acquiring mass spectrometry data. The mass spectrometer was purchased via an NIH-shared instrument grant, S10 OD023436. This work was supported by grants from the National Eye Institute (R01EY029680, R01EY028884, and R01EY036086 to Y.I.). This work was partly supported by a Challenge Grant from Research to Prevent Blindness to the Department of Ophthalmology, Indiana University School of Medicine, and by an award from the Ralph W. and Grace M. Showalter Research Trust and the Indiana University School of Medicine to Y.I. This work was also partly supported by Cohen Pilot Grants in Macular Degeneration Research (to Y.I.) from the Department of Ophthalmology, Indiana University School of Medicine.

## Author contributions

S.T. and Y.I. designed the research project. S.T. conducted the experiments and analyzed the data presented in Figures 1, 2, 3, 4, 5, and 8. H.H. conducted the experiments and analyzed the data presented in Figures 6 and 7. M.M. contributed to the acquisition and analysis of the mass spectrometry data. Y.I. wrote the entire manuscript. S.T. wrote the result and method sections, wrote the draft, and edited the entire manuscript. H.H. and M.M. edited the manuscript. All the authors approved the final version of the manuscript.

## Data availability

The raw RNA-seq data used in this study have been deposited to the NCBI. All identified RNAs are listed in Supplemental Table S1.

---

**Supplemental Figure S1.**
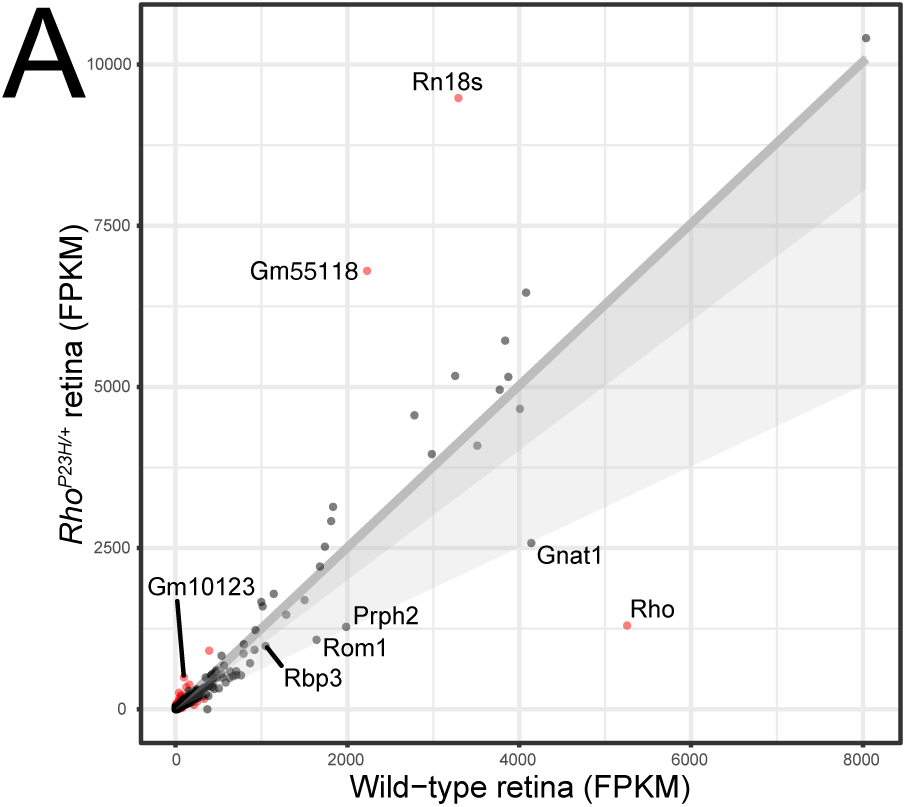
*Rho* mRNA is downregulated in P35 *Rho^P23H/+^* retinas. (A) RNA expression levels were plotted to compare P35 wild-type and *Rho^P23H/+^* mouse retinas reared under standard cyclic 200 lux conditions. A total of 811 genes (shown in red dots) are differentially expressed with 2-fold change (log2-transformed change of < -1 or > 1) and q-value of < 0.05 thresholds. *Rho* was the second most abundantly expressed gene in wild-type retinas, and was downregulated by 75.3% in *Rho^P23H/+^* retinas. The dark gray shade indicates less than a 20% decline in mRNA expressions. The light gray shade indicates less than a 50% decline in mRNA expressions.

**Supplemental Figure S2.**
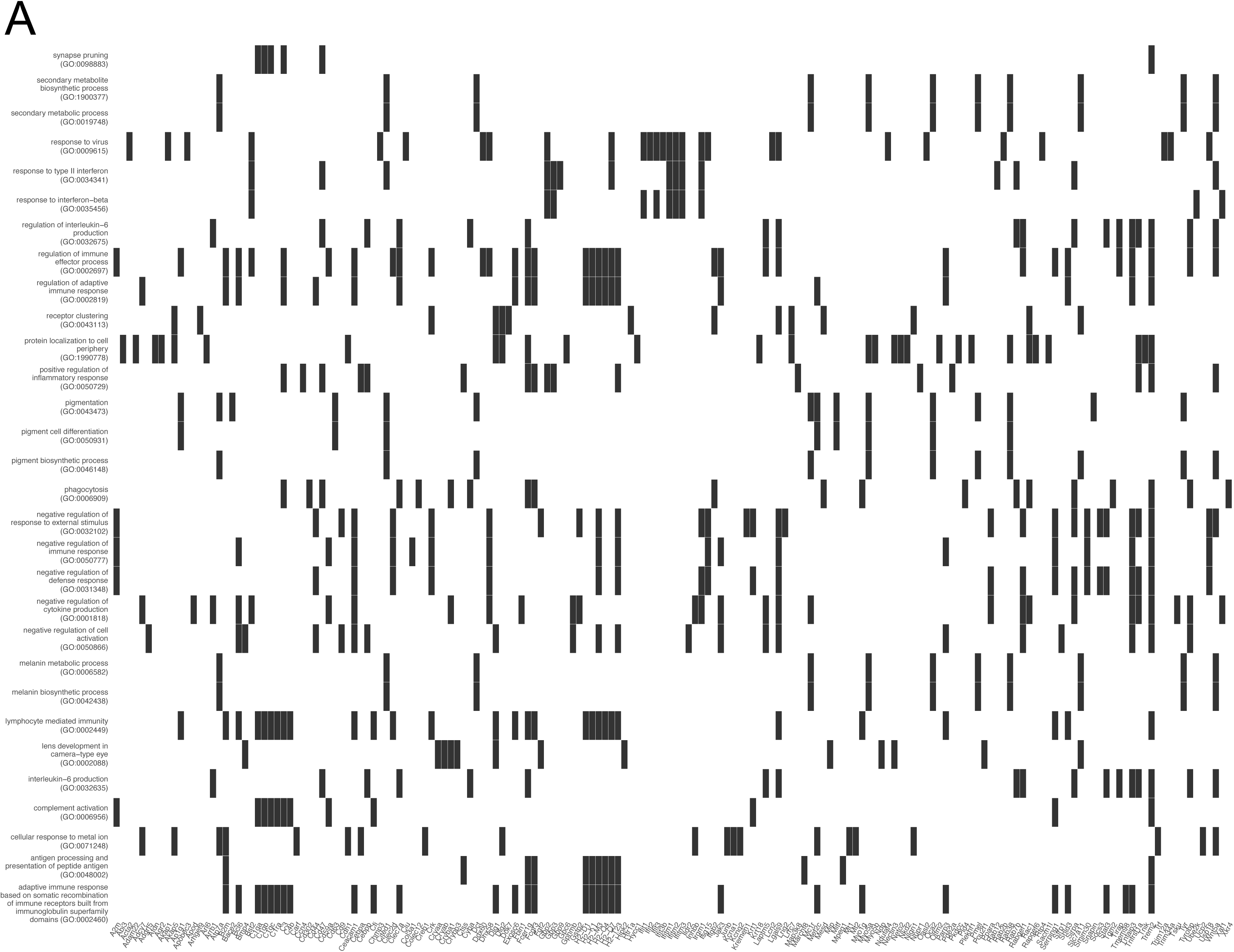
Associations between Gene Ontology (GO) terms for biological processes and specific genes for P35 *Rho^Q344X/+^* retinas in comparison to wild-type retinas under standard cyclic 200 lux light condition. (A) Gene names are shown on the x-axis and significantly enriched GO terms are shown on the y-axis. Black rectangles show their corresponding matches and visualize the associations between GO terms and protein names.

**Supplemental Figure S3.**
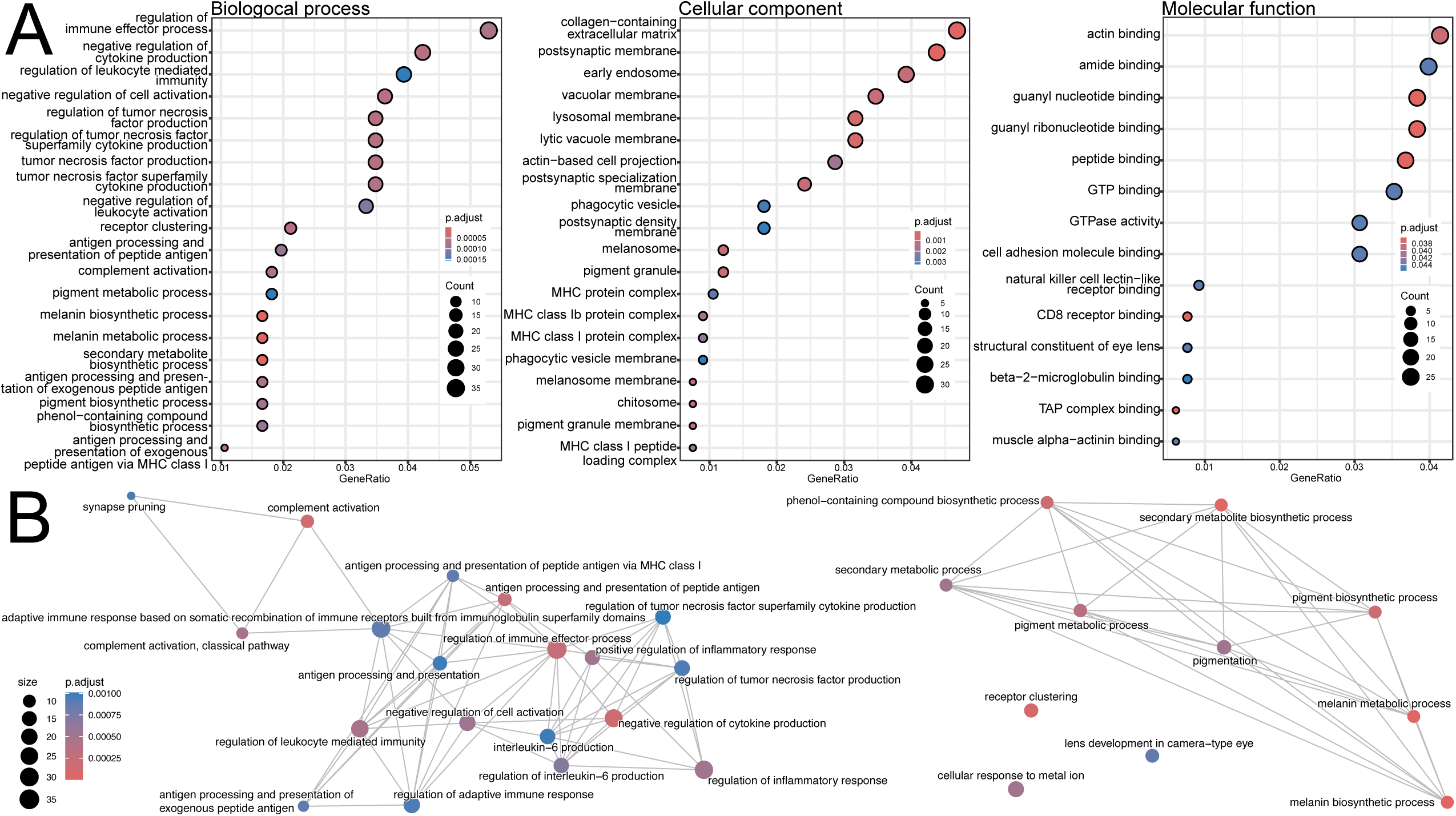
Comparison of RNA-seq between P35 wild-type and *Rho^Q344X/+^* retinas identifies that inflammation-related genes are upregulated in *Rho^Q344X/+^*retinas. (A) Pathway analyses for GO terms using the mRNA expression with 2-fold change are shown. Significantly changed pathways (up to 20) are shown for biological process, cellular component, and molecular function. The size of each dot within the plots represents the level of protein enrichment (Count), and the color coding indicates statistical significance (p.adjust), as shown on the right side of each panel. (B) Networks illustrating the output of the hypergeometric test. The pathway analysis was based on gene enrichment. The size of each dot (Count) represents the level of gene enrichment (number of genes in each category), and the color coding shows statistical significance (p.adjust), as indicated at the bottom left corner.

**Supplemental Figure S4.**
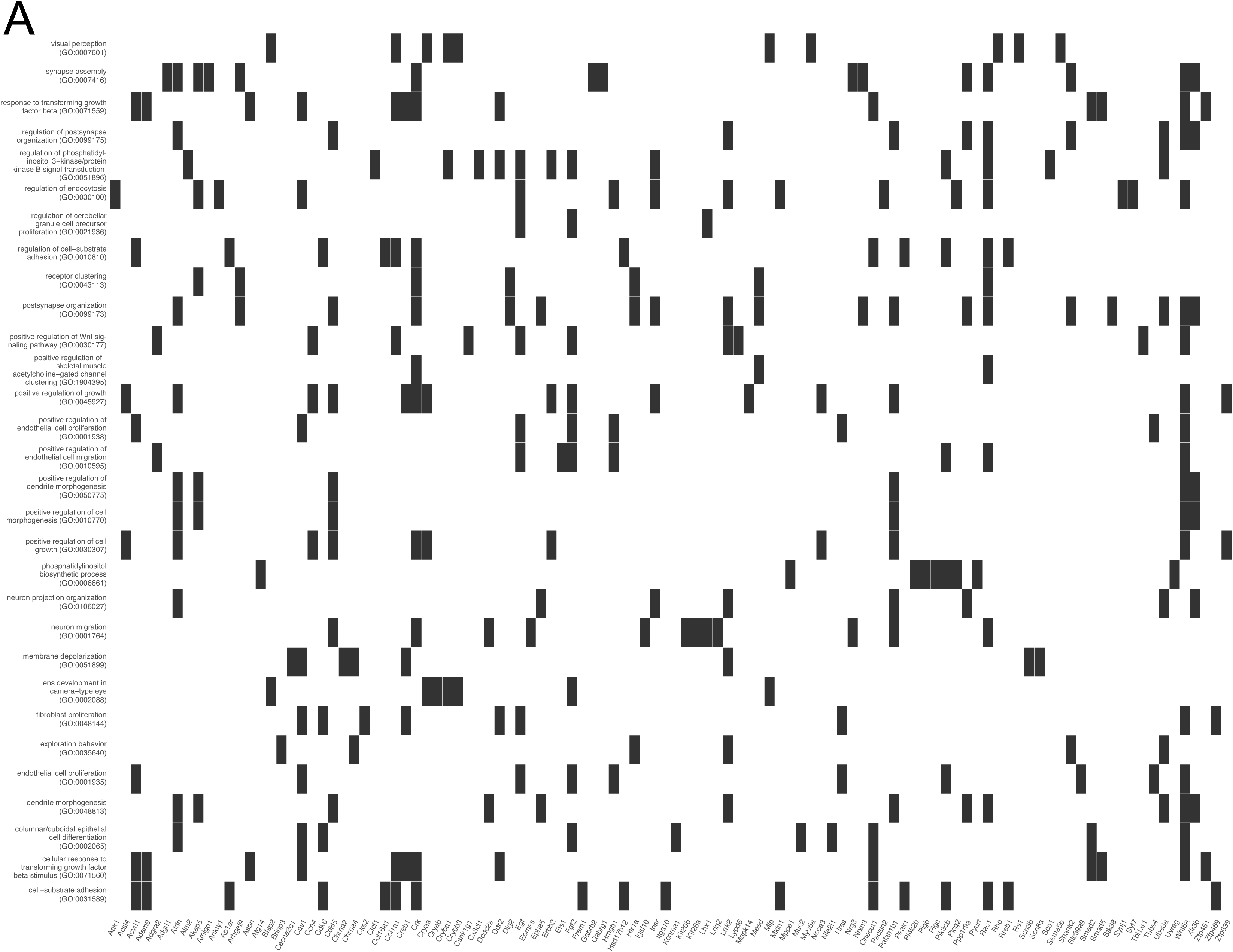
Associations between GO terms for biological processes and specific genes for P35 wild-type retinas reared under dark condition. (A) Gene names are shown on the x-axis and significantly enriched GO terms are shown on the y-axis. Black rectangles show their corresponding matches and visualize the associations of enriched GO terms and protein names.

**Supplemental Figure S5.**
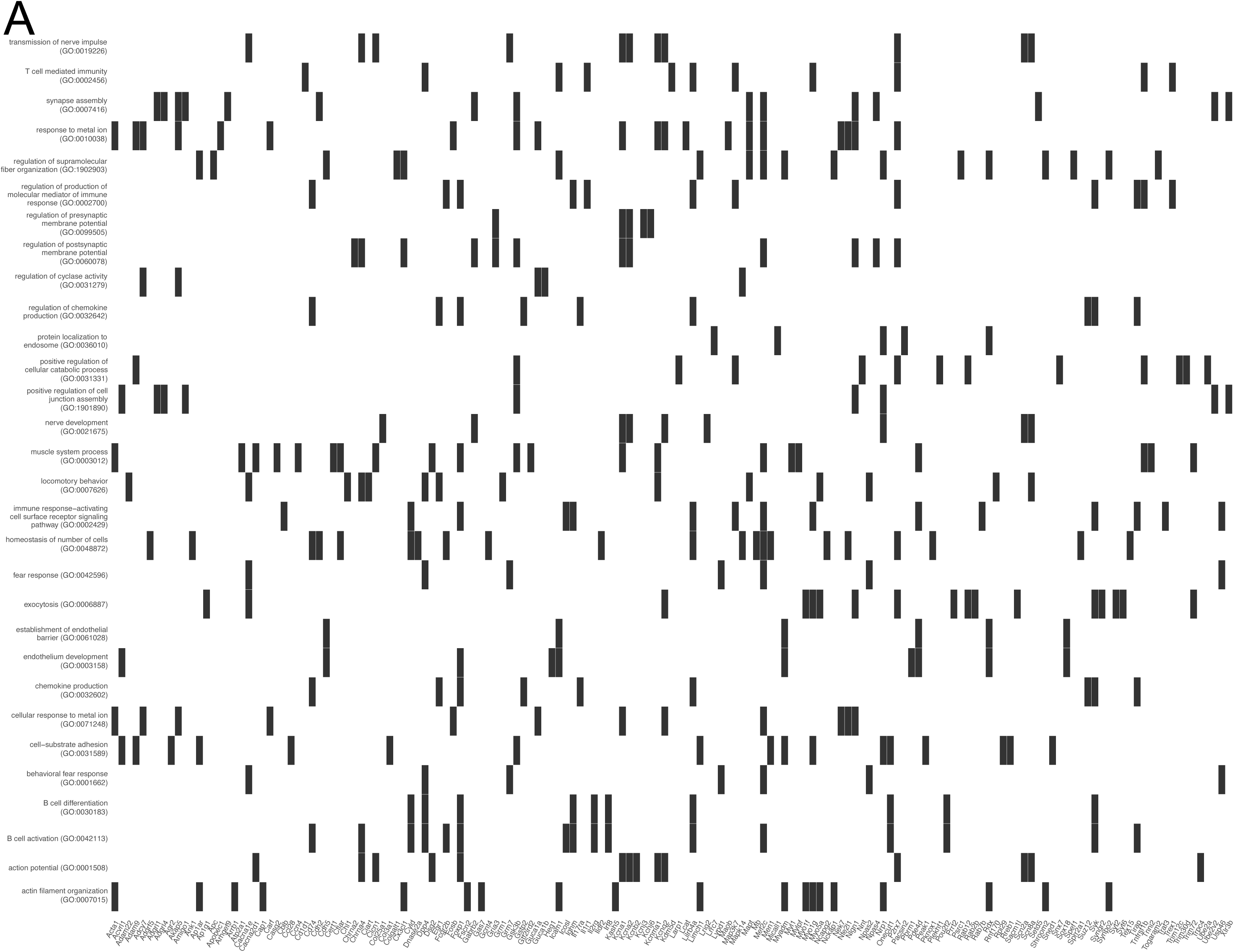
Associations between GO terms for biological processes and specific genes for P35 *Rho^Q344X/+^*retinas reared under dark condition. (A) Gene names are shown on the x-axis and significantly enriched GO terms are shown on the y-axis. Black rectangles show their corresponding matches and visualize the associations of enriched GO terms and protein names.

